# Defining bottlenecks and opportunities for Lassa virus neutralization by structural profiling of vaccine-induced polyclonal antibody responses

**DOI:** 10.1101/2023.12.21.572918

**Authors:** Philip J.M. Brouwer, Hailee R. Perrett, Tim Beaumont, Haye Nijhuis, Sabine Kruijer, Judith A. Burger, Wen-Hsin Lee, Helena Müller-Kraüter, Rogier W. Sanders, Thomas Strecker, Marit J. van Gils, Andrew B. Ward

## Abstract

Lassa fever continues to be a major public health burden in endemic countries in West Africa, yet effective therapies or vaccines are lacking. The isolation of potent and protective neutralizing antibodies against the Lassa virus glycoprotein complex (GPC) justifies the development of vaccines that can elicit strong neutralizing antibody responses. However, Lassa vaccines candidates have generally been unsuccessful in doing so and the associated antibody responses to these vaccines remain poorly characterized. Here, we establish an electron-microscopy based epitope mapping pipeline that enables high-resolution structural characterization of polyclonal antibodies to GPC. By applying this method to rabbits vaccinated with a recombinant GPC vaccine and a GPC-derived virus-like particle, we reveal determinants of neutralization which involve epitopes of the GPC-C, GPC-A, and GP1-A competition clusters. Furthermore, by identifying previously undescribed immunogenic off-target epitopes, we expose challenges that recombinant GPC vaccines face. By enabling detailed polyclonal antibody characterization, our work ushers in a next generation of more rational Lassa vaccine design.

## Introduction

Lassa fever, a viral hemorrhagic fever which is caused by the Lassa virus (LASV), is a persistent public health and socioeconomic burden to affected countries in West Africa. LASV is predominantly transmitted to humans from its natural reservoir *Mastomys natalensis*^1^(though person-to-person transmission does occur^2–4^), infecting an estimated 500,000-900,000 people each year and resulting in approximately 5,000 deaths.^5,6^ While many people infected with LASV have asymptomatic or mild infections, those that present with clinical symptoms suffer from fever, encephalitis, respiratory distress, facial swelling, bleeding from orifices, and multiorgan failure, which without early treatment may result in death.^7^ Even in cases of mild Lassa fever, approximately one-third of those infected experience sensorineural hearing loss, which results in socioeconomic strain in endemic regions.^8–10^ Further, current estimates suggest those affected by the disease while pregnant have almost three-fold higher fatality rates and 80% of these patients experience intrauterine fetal deaths.^11^ With no effective and approved therapeutic or vaccine, the World Health Organization has classified Lassa fever as a priority disease.^12^ The development of a vaccine that can protect against this devastating pathogen would be a major public health advancement.

Lassa vaccine efforts focus primarily on the Lassa virus glycoprotein complex (GPC). Not only does GPC harbor numerous T-cell epitopes,^13,14^ it is also the sole mediator for host cell infection and presents all known neutralizing antibody (NAb) epitopes.^15^ The GPC resides on the viral surface as a trimer of GP heterotrimers with each GP protomer comprised of the receptor-binding subunit GP1; the transmembrane-spanning subunit GP2; and the stable signal peptide (SSP), which remains non-covalently attached to GP after it is cleaved from the GP precursor.^16–19^ The prefusion GPC facilitates infection of host cells by engaging with its primary extracellular receptor matriglycan.^20–22^ After internalization and trafficking to the endosome via macropinocytosis,^23^ GP1 undergoes a conformational change in response to the endosome’s acidic pH, which enables its binding to the endosomal receptor lysosomal-associated membrane protein 1 (LAMP-1).^24,25^ LAMP-1 binding facilitates GPC’s conformational shift from a pre- to post-fusion state and promotes viral and host membrane fusion.^26^ The conformational lability of GPC, required to undergo these dramatic structural changes, has frustrated the development of stable recombinant prefusion GPC for many years. The introduction of prefusion-stabilizing GPCysR4 mutations^27^—R207C and G360C to introduce a disulfide bond between GP1 and GP2, the helix-breaking E329P, and L258R and L259R to replace the site-1 protease (S1P) cleavage site with a furin cleavage site—combined with the complexing of a trimer-stabilizing monoclonal antibody (mAb) 37.7H finally enabled the generation and high-resolution structural characterization of prefusion GPC trimers.^27,28^ Since then, methods have been described to stabilize the trimeric conformation of prefusion GPC in the absence of a trimer-stabilizing mAb.^29–31^ We previously demonstrated that prefusion GPC trimers are stabilized by genetic fusion to the trimeric scaffold I53-50A.^30,31^ These fusion proteins, known as GPC-I53-50A, have not only proven to be a useful reagent for characterizing and isolating GPC-specific antibodies but also were able to induce NAb responses in rabbits.^30,31^

Protective immunity to LASV infection has been attributed to cell-mediated responses^5^ with severe infections often associated with poor T-cell responses to the GPC and nucleoprotein.^14,32,33^ NAb responses appear to play a limited role in curbing acute infection as they generally appear months after viral clearance.^34^ The GPC’s dense glycan shield and the elicitation of non-NAbs against the post-fusion or uncleaved conformations of the GPC likely hinder the induction of NAbs.^35–38^ In addition, by a mechanism that remains unknown, LASV delays IgM class-switching to IgG, further impeding the maturation of NAb responses.^39^ Nevertheless, numerous potent NAbs have been isolated from convalescent individuals and the mechanism of action for several of these NAbs has been elucidated.^15,27–29,31,40,41^ Cocktails of isolated NAbs have shown great promise as therapeutics in animal models. For example, Arevirumab-3, a cocktail of NAbs 8.9F, 37.2D and 12.1F, shows protection in non-human primates after LASV challenge.^42,43^ These early successes justify pursuing the development of vaccines that are capable of inducing potent NAb responses.

Recent efforts in Lassa vaccine design include an mRNA vaccine; DNA-based vaccine^44^; measles, rabies, adenovirus, and vesicular stomatitis virus (VSV) based vector vaccines^45–50^; nanoparticle-based protein vaccine^30^; and virus-like particle vaccines.^51,52^ While many of these have shown efficacy in animal models, induction of NAbs in most cases was absent or required numerous boosting immunizations. The reasons for this may vary between vaccine platforms but may include possible presentation of irrelevant GPC conformations to B cells. Nevertheless, careful and robust assessments of vaccine-induced antibody responses have been lacking and, as a result, it remains unclear which epitopes are being targeted on this highly glycosylated glycoprotein. Identifying the vaccine-induced on- and off-target responses and understanding their place in the immunodominance hierarchy will be an important first step in designing next-generation LASV vaccines that induce more potent NAb responses.

In addition to traditional assays to assess humoral immune responses, electron-microscopy based polyclonal epitope mapping (EMPEM) has been used successfully to add low- and/or high-resolution structural definition to polyclonal antibody (pAb) immune responses across several pathogens.^53–57^ In short, serum antibodies from immunized or challenged individuals can be purified, digested to Fab, complexed with antigen of interest, and directly visualized using electron microscopy (EM). This methodology is valuable for determining the presence and immunodominance of responses across time. Here, we describe the development and in-depth characterization of a next-generation GPC-I53-50A trimer and use it to establish an EMPEM pipeline for LASV. We then apply this method to investigate the pAb responses from rabbits previously immunized with GPC-I53-50A as well as a GPC-derived VLP.^30,52^ Our analysis identified a range of epitopes that were targeted by pAbs, including neutralizing epitopes of the GPC-C and GPC-A competition groups. In addition, we describe off-target, non-neutralizing epitopes at the interior and base of the recombinant GPC trimer. By generating high-resolution models of the epitope-paratope interfaces of vaccine-induced Ab responses we provide a detailed landscape of the immunogenic sites on GPC in different vaccine contexts. Our EMPEM work not only reveals the challenges that recombinant LASV vaccines face in inducing NAb responses but also informs the design of the next iteration that may overcome them.

## Results

### Development and characterization of recombinant LASV GPC trimers that bind 8.9F

We recently showed that fusing GPCysR4 to I53-50A resulted in the generation of prefusion GPC trimers that present epitopes of the GP1-A, GPC-A, and GPC-B competition clusters; however, these immunogens still lacked the ability to bind 8.9F mAb, the sole known member of the GPC-C cluster. This highly potent broadly neutralizing antibody (bNAb) directly blocks matriglycan binding and has been shown to require the native cleavage site to engage with GPC.^29^ Therefore, to generate a trimeric GPC immunogen that presents all currently known bNAb epitopes, we reverted the RRRR site of GPCysR4 to the native RRLL sequence. The resulting construct, GPCysRRLL-I53-50A, was expressed in HEK 293F cells by co-transfection with S1P and purified by subsequent Strep-Tactin XT affinity chromatography and size-exclusion chromatography (SEC) steps (Figure 1A). Sodium dodecyl sulfate polyacrylamide gel electrophoresis (SDS-PAGE) demonstrated these proteins were efficiently cleaved (Figure S1A) while negative stain electron microscopy (nsEM) confirmed that GPCysRRLL-I53-50A forms trimers (Figure 1B). Furthermore, GPCysRRLL-I53-50A showed binding to representative bNAbs of the GP1-A (12.1F), GPC-A (25.10C) and GPC-B (37.7H) epitope cluster (Figure S1B), while restoring its ability to bind 8.9F (Figure 1C).

**Figure 1.**
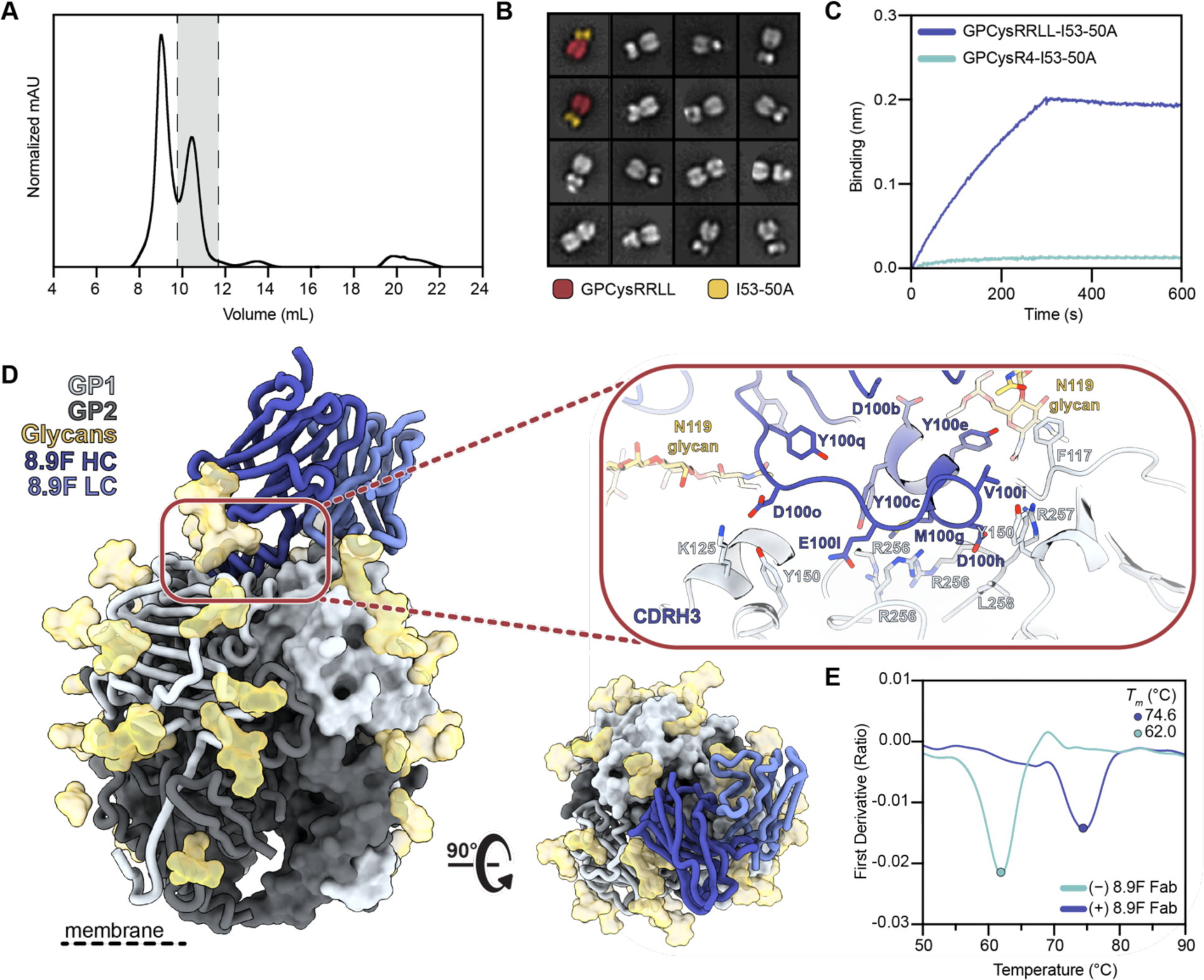
Development and characterization of GPCysRRLL-I53-50A and binding to 8.9F. **(A)** Representative SEC of GPCysRRLL-I53-50A. Fractions containing trimer are shown in grey. **(B)** 2D class averages from nsEM of the GPCysRRLL-I53-50A. The far left GPCysRRLL-I53-50As are pseudo-colored and show the GPCysRRLL (red) and I53-50A scaffold (gold). **(C)** Representative BLI sensorgram comparing binding of immobilized 8.9F IgG to GPCysRRLL-I53-50A (blue) and GPCysR4-I53-50A (cyan). Single curves are shown that are representative of three technical replicates. **(D)** Atomic model of GPCysRRLL-I53-50A bound to 8.9F Fab (blue) determined by cryo-EM. Inset depicts key interactions between GP1 and 8.9F at the epitope-paratope interface. Glycan N119, which makes extensive contacts with the heavy and light chains, is shown in gold. More details on the epitope-paratope interactions are presented in Table S1. **(E)** Thermostability of GPCysRRLL-I53-50A with and without 8.9F Fab bound assessed by nanoDSF. Melting temperatures (*T*_m_) are calculated as the inflection point (circles) of the ratio of signal at 350 and 330 nM. Each melting curve is a representative of triplicate curves with *T*_m_ within ±0.1°C.

Next, to structurally characterize GPCysRRLL-I53-50A in molecular detail, we used single-particle cryogenic electron microscopy (cryo-EM) to solve a 3.0 Å structure of GPCysRRLL-I53-50A in complex with 8.9F Fab (Figure 1D). Our GPC model shows a near-identical structural homology to that of a previously described detergent-solubilized full-length native GPC complexed with 37.2D and 8.9F Fab, as measured by the root-mean square deviation (RMSD; 0.53 Å among the full 8.9F sequence and 190 pruned atom pairs per GP1; 1.0 Å overall fit across all sequence-aligned pairs with PDB 7UOT^29^; Figure S1C). The most notable difference between the models was at residues 204–214 in GP1 where the presence of the introduced prefusion-stabilizing disulfide bond in GPCysRRLL-I53-50A leads to an alternative conformation of this flexible loop. Consistent with earlier observations, 8.9F targets the trimer apex, making contacts with all three protomers while using its 31-amino acid CDRH3 to penetrate the three-fold axis in between the α1 helix and the complex-type glycans at position N119 (Table S1). Meanwhile the LC interacts with the neighboring protomer, principally making contacts with the N119 glycan. Considering the way 8.9F engages with the RRLL sequence on all three protomers of GPC, we hypothesized that 8.9F may have a stabilizing effect on recombinant GPC trimers. Indeed, nano differential scanning fluorimetry (nanoDSF) studies showed that the thermostability of GPCysRRLL-I53-50A increases by 12°C when complexed with 8.9F (*T_m_*=62°C for GPCysRRLL-I53-50A; *T_m_*=74°C for GPCysRRLL-I53-50A-8.9F Fab; Figure 1E). Interestingly, in contrast to earlier suggestions that the leucines of the S1P cleavage site stabilize the trimeric interface,^16^ we observed a slightly lower thermostability of GPCysRRLL-I53-50A than GPCysR4-I53-50A (Figure S1D). Overall, we conclude that we have developed a new generation of prefusion-stabilized trimeric GPC-I53-50A with an improved antigenic profile.

### Development of an EMPEM protocol for LASV GPC trimers

EMPEM is a powerful method to map antigen-specific pAb responses induced by infection or vaccination.^53^ While EMPEM studies of HIV-1,^53,54,57–61^ influenza,^56,62^ and coronavirus^55^ vaccine-induced antibody responses has boosted further vaccine development, the antibody profile induced by Lassa vaccines remains ill-defined. For our initial attempts to map GPC-specific polyclonal responses we used pools of purified Fabs from rabbits that were previously immunized with GPCysR4-I53-50A trimers.^30^ Overnight incubations of GPCysR4-I53-50A with an excess of rabbit Fabs resulted in the formation of immune complexes as shown by the shift in the SEC trace of GPCysR4-I53-50A. However, when the corresponding fractions were pooled, imaged by nsEM, and subjected to single-particle extraction and 2D classification, we only observed classes of GP monomers (i.e. splayed-open GPC held together by the I53-50A scaffold) were discernible (data not shown). To assess if this issue could be mitigated by incubating for a shorter time-period, we generated complexes and incubated for 4 hours. Again, 2D class averaging of extracted single-particles did not generate classes that represented trimeric GPC-Fab complexes (Figure 2A, top right). These results seemed to reveal that pAbs cause disassembly of our recombinant GPC trimers.

**Figure 2.**
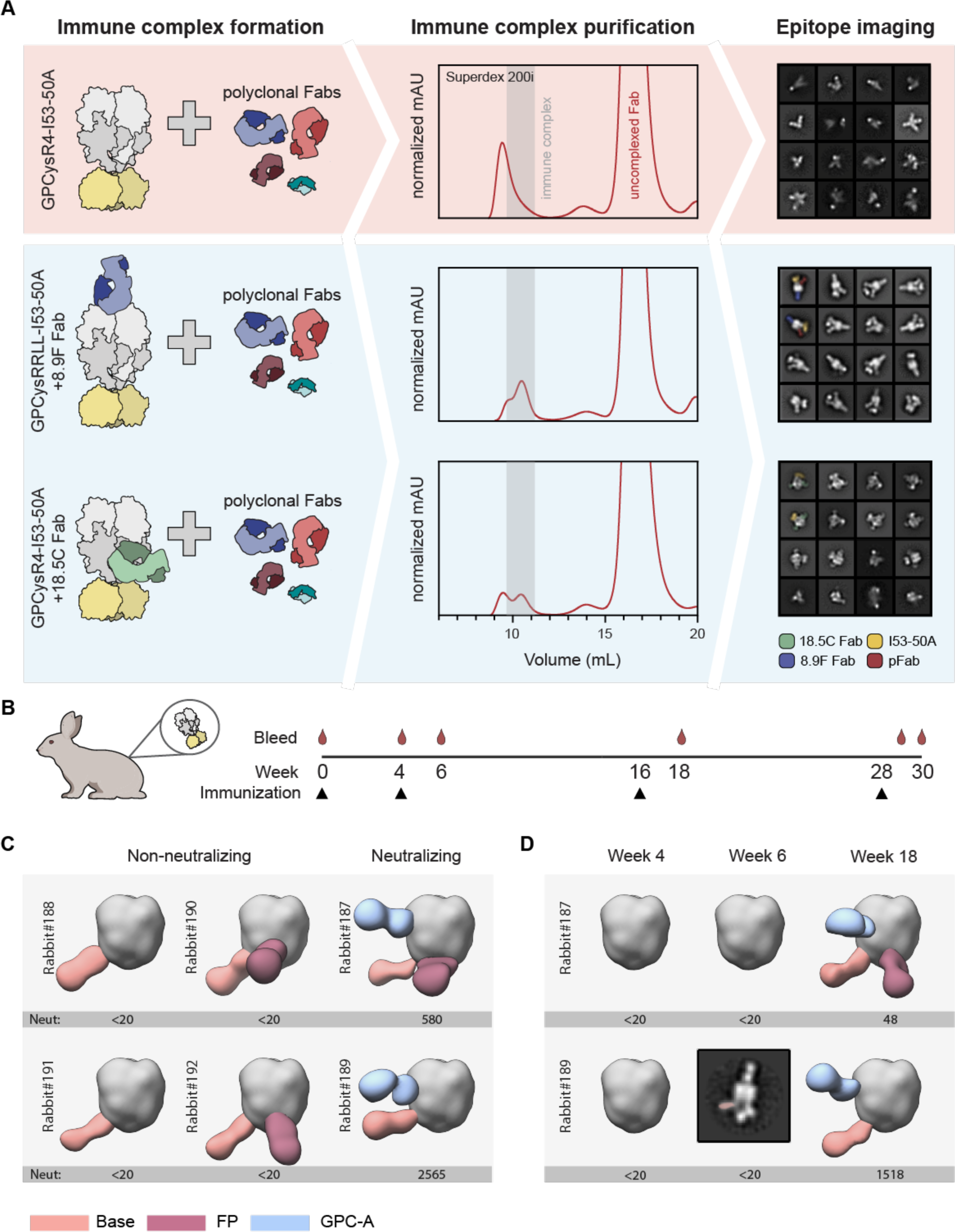
Stabilizing antibodies facilitate LASV GPC EMPEM and generation of epitope landscapes. **(A)** Schematic of LASV GPC EMPEM development. In the absence of stabilizing antibodies, GPC undergoes antibody-facilitated disassembly, which leads to aggregation as determined by SEC and splayed open GPC trimers as visualized by nsEM (red box). In the presence of stabilizing Fabs 8.9F and 18.5C, immune complexes are formed and the pAb response against GPC trimers can be visualized enabling mapping of the pAb landscape (blue box). **(B)** Immunization scheme of the six rabbits immunized at week 0, 4, 16 and 28 with GPCysR4-I53-50A antigen, as recently described.^30^ **(C-D)** Composite figures of pAb responses of each individual rabbit as determined by nsEMPEM. To the left of each figure the rabbit identifier is shown and the previously described ID_50_ of LASV pseudovirus serum neutralization is depicted below.^30^ A titer of <20 represents no neutralization. pAbs are color coded based on their apparent epitope as shown in the legend. For simplicity only a single Fab is shown and consistently projected on the same protomer. **(C)** Binding of pAbs at week 30. **(D)** Binding of pAbs at week 4, 6, and 18 for rabbit 187 and 189. Because of limited numbers of particles that contain a base response, we were unable to reconstruct a 3D map for the week 6 pAb response in rabbit 189. Instead, a 2D class is shown with the base pAb highlighted by pseudocoloring.

Considering 8.9F’s effect on the thermostability of GPC trimers, we rationalized that complexes of 8.9F Fab with GPCysRRLL-I53-50A may be able to withstand the Fab-induced disassembly and prove to be a useful reagent for EMPEM studies. Indeed, when we performed single-particle analysis from nsEM studies with GPCysRRLL-I53-50A-8.9F complexed with polyclonal rabbit Fabs, we were able to obtain a sufficient amount of 2D class averages that represented Fab-bound GPC trimers (Figure 2A, middle right). As using GPCysRRLL-I53-50A-8.9F complexes to probe pAb responses to GPC occludes potential apex-targeting pAbs, we also performed EMPEM with the same polyclonal rabbits Fabs using GPCysR4-I53-50A complexed with the trimer-stabilizing Fab 18.5C (Figure 2A, bottom). Here, GPCysR4-I53-50A was chosen instead of GPCysRRLL-I53-50A so that it matched the vaccine sequence. Although in this context pAb binding was not observed, the trimers clearly withstood pAb-induced disassembly. We thus present a two-pronged approach for LASV EMPEM: (1) pre-complexing GPCysRRLL-I53-50A with 8.9F Fab to sample pAb responses excluding those targeting the apex, and (2) pre-complexing with 18.5C Fab to sample pAbs which target the apex (Figure 2A).

### Analysis of recombinant LASV GPC-induced antibody responses: the base epitope

Having established an EMPEM protocol, we continued to apply it to all six rabbits that were previously immunized with GPCysR4-I53-50A.^30^ These rabbits received an intramuscular immunization of GPCysR4-I53-50A formulated in Squalene Emulsion adjuvant at weeks 0, 4, 16, and 28 (Figure 2B).^30^ Two out of these rabbits induced pseudovirus neutralization titers and four rabbits did not (Figure 2C), which enabled us to correlate neutralization with mapped epitopes. The resulting 3D representations of GPC with bound pAbs from week 30 revealed a remarkably consistent response targeting the GPC base (Figure 2C, Figure S2A and Table S2). These antibodies were present in all rabbits, regardless of whether they induced neutralizing responses, and thus may represent a common but non-neutralizing response to recombinant GPC immunogens. To understand the evolution of these responses during the prime-boosting strategy we performed EMPEM experiments with week 4, 6 and 18 serum samples from rabbits 187 and 189 (Figure 2D, Figure S2B, and Table S2). Base-targeting responses were the only responses discernible at week 6 (after secondary immunization), albeit at very low levels and only in one out of two rabbits, highlighting the relative immunogenicity of this epitope.

To obtain more detailed structural information on the epitope-paratope interface of these base-targeting responses, we performed single-particle cryo-EMPEM on GPC-Fab complexes from rabbit 190. We reconstructed two high-resolution maps of structurally unique GPC-Fab complexes with each map revealing a Fab targeting the GPC base (Figures S3 and S4, Table S3). Of the two maps, corresponding to Base-1 and Base-2, the former had sufficient resolution to guide initial placement of conserved amino acids. This enabled us to relax previously deposited rabbit Fab models and define CDR loops with greater confidence.^30,63^ After initial placement and relaxation, the models’ amino acids were re-defined as poly-alanine to account for our lack of sequence information (Figure 3A). Our model showed that Base-1 targets the N-terminal loop of GP1 (aa 59–67) and several residues at the beginning of the C-terminal helix of GP2 (aa 395–419). The N-terminal loop is engaged by the pAbs’ CDRH1, CDRH2, CDRH3, and CDRL1 whereas the C-terminal helix is targeted using its CDRL1 and CDRL2. Furthermore, Base-1 seemed to make contacts with both N373 and N395 glycans. Despite insufficient resolution of the Base-2 map for model building, a docked rabbit Fab model (PDB: 7RA7) in the pFab density revealed a clearly distinct angle of approach compared to Base-1 (Figure S5A). With respect to the C-terminal helix, Base-1 approaches GPC in a near-horizontal plane while Base-2 is tilted approximately 80° and binds at a much steeper angle (Figure S5B). These differences imply that the two Fabs originate from different germline B cells, which would be in line with the observed immunogenicity of this epitope.

**Figure 3.**
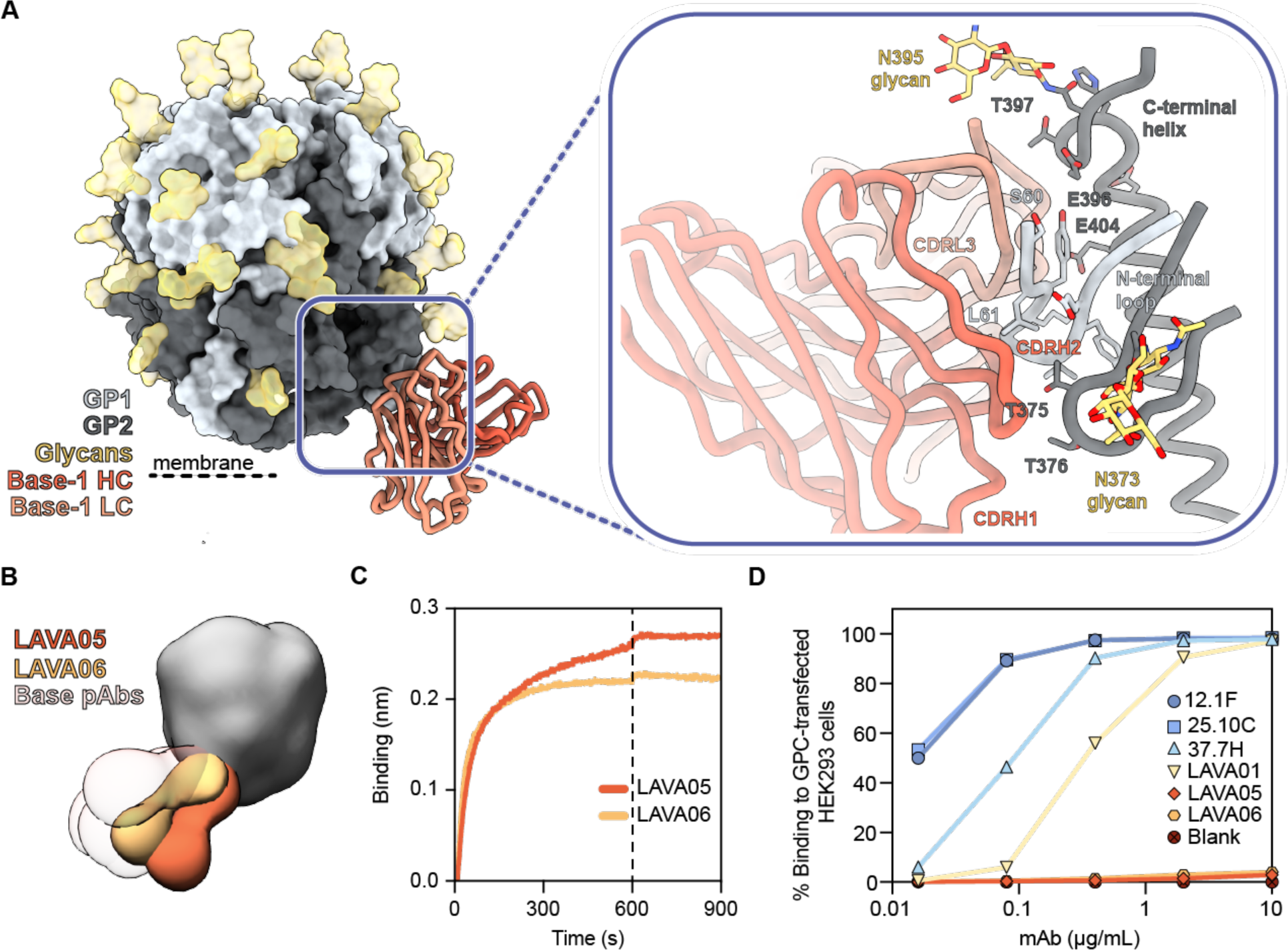
Recombinant LASV GPC immunization in rabbits induces antibody responses to the GPC base. **(A)** Atomic model of Base-1 Fab bound to GPCysRRLL-I53-50A determined by cryo-EM. Inset depicts presumed interactions between the modeled polyalanine Fab model and the GPC. **(B)** Composite figure of LAVA05 and LAVA06 binding to GPC as determined by nsEM. Base-targeting pAb responses from rabbits 187-192 are presented with transparent coloring to display the overlap between the mAbs and pAbs. **(C)** Sensorgrams from BLI experiments measuring binding of LAVA05 and LAVA06 to GPCysRRLL-I53-50A captured by his-tagged 8.9F Fab immobilized on Ni-NTA biosensors. **(D)** Binding of mAbs to full-length native GPC expressed on HEK293 cells. MAbs bound to GPC-expressing cells were detected by flow-cytometry using PE-labeled anti-rabbit/human IgG. Percentage of fluorescently labeled cells are depicted as a function of mAb concentration.

Superimposition of the models of Base-1 with a model of a full-length GPC (PDB: 7PUY^16^) suggested that this base-targeting response may not be elicited when GPC is embedded in the membrane. This response would likely be blocked by the lipid bilayer in the context of native virions. Additionally, if the membrane-embedded GPC were able to tilt, binding of this Fab to the base epitope could be sterically hindered by the non-covalently bound SSP (Figure S5C). To investigate if the base epitope is accessible on membrane-embedded GPC, we isolated two mAbs which shared the same HC sequence: LAVA05 and LAVA06. These mAbs bound GPC in a very similar fashion as the base-targeting responses observed in rabbit sera (Figure 3B, Figure S5D and Table S2). Despite their strong binding to GPC (Figure 3C), LAVA05 and LAVA06 did not neutralize LASV pseudovirus (Figure S5E), consistent with our EMPEM observations that base-responses do not correlate with neutralization (Figure 2C). Furthermore, while rabbit NAb LAVA01 and human NAbs 12.1F, 25.10C, and 37.7H showed binding to GPC embedded on the membrane of transfected HEK293 cells, LAVA05 and LAVA06 did not (Figure 3D). This data, together with our EMPEM structures, suggest that the base constitutes a highly immunogenic neo-epitope that is made accessible when generating recombinant GPC trimers.

### Analysis of recombinant LASV GPC-induced antibody responses: the GPC-A epitope

Beyond the omnipresent base responses, Fab densities targeting an equatorial epitope were noticeable from the week 30 negative-stain EMPEM data (Figure 2C). Interestingly, these responses were only present in the two rabbits that generated NAbs (rabbit 187 and 189) suggesting that these pAbs might be neutralizing (Figure 2C). Longitudinal EMPEM analyses with serum from weeks 4, 6 and 18 revealed Abs against these equatorial epitopes are only being elicited after three immunizations at week 18 (Figure 2D). Interestingly, week 18 is also when we see the emergence of pseudovirus NAb titers, further strengthening the correlation between these equatorial responses and neutralization.

High-resolution structural information on the interaction between these equatorial responses and GPC was obtained by performing cryo-EM with immune complexes of week 30 serum from rabbit 187 (Figures S3 and S4, Table S3). Our data reveals the equatorial-targeting pAb engages the GPC in a manner reminiscent of GPC-A-specific NAbs 36.1F, and, to a lesser extent, 25.10C (Figures 4A and 4B, Figures S6A).^40^ This pAb, which we named GPC-A-1, predominantly interacts with one GP1 protomer where it uses its 13 amino acid CDRH3 loop to make contact with the flexible loop that extends across the beta-sheet surface near the GP1-GP2 interface. The CDRH3 engages residue E76 between η1 and β2 in a manner very similar to 36.1F’s CDRH3.^40^ Furthermore, our model suggests that GPC-A-1 engages with all contacted residues shared by the GPC-A-targeting Abs 36.1F and 25.10C: N74, E76, K88, T226, W227, E228, D229, H230, and Q232 (Figure 4A).^40^ Typical of GPC-A antibodies, GPC-A-1 extensively targets glycans and uses its CDRL3 and CDLR1 loops to make contact with glycans N365 and its CDRH2 loop to interact with N89. Glycans N224 and N79 surround the GPC-A-1 HC and LC, respectively, though we hypothesize their role at the epitope is modest compared to N365 and N89. Overall, high-resolution characterization of the epitope-paratope interface of GPC-A-1 revealed a remarkable similarity to known NAbs, which, together with the strong correlation between neutralization and the presence of GPC-A-1-like pAbs, implies that these responses were the driving force behind LASV neutralization in these rabbits.

**Figure 4.**
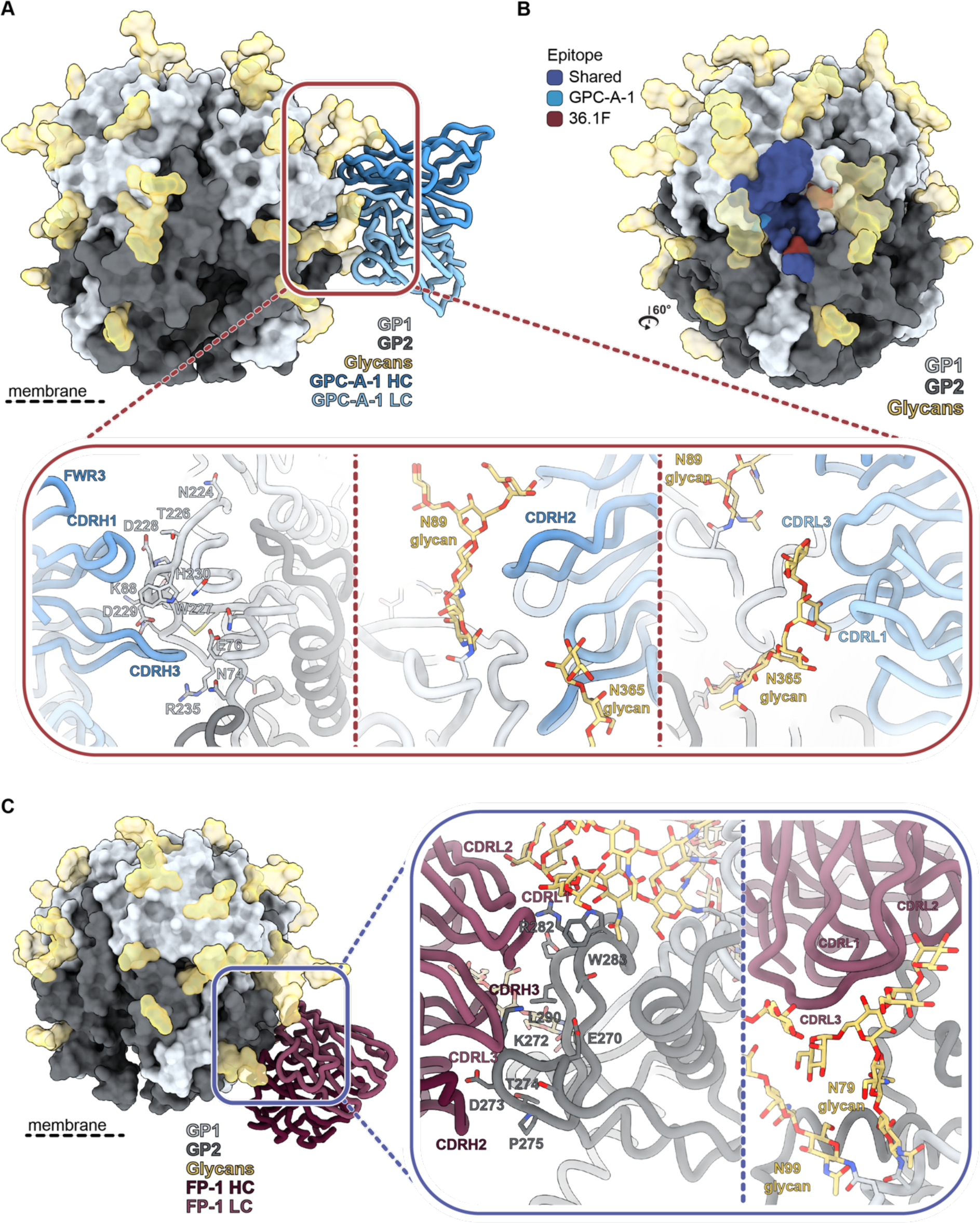
Recombinant LASV GPC immunization in rabbits induces antibody responses to the GPC-A and fusion peptide epitope. **(A)** Atomic model of GPC-A-1 Fab bound to GPCysRRLL-I53-50A determined by cryo-EM. Inset depicts putative interactions on GPC at the epitope-paratope interface. Important residues shared by GPC-A antibodies 36.1F and 25.10C are shown as sticks.^40^ **(B)** Epitope footprint comparison between GPC-A-1 and 36.1F Abs. Epitope-Analyzer^95^ was used to determine (putative) contact residues using a cut-off of 4 Å between epitope and paratope for 36.1F and 6 Å for GPC-A-1 (poly-alanine model). **(C)** Atomic model of FP-1 Fab bound to GPCysRRLL-I53-50A as determined by cryo-EM. Inset depicts residues of GPC, shown as sticks, that make putative contact with the polyalanine Fab.

### Analysis of recombinant LASV GPC-induced antibody responses: the fusion peptide epitope

Cryo-EM analysis of the polyclonal response in rabbit 190 enabled us to provide high-resolution detail of a response observed in 3 out of 6 rabbits which targeted an epitope in close proximity to the base epitope described earlier (Figures 2C, Figures S3 and S4, and Table S3). Our model revealed an antibody targeting the fusion peptide, which we refer to as FP-1. This pAb binds to GPC in a loosely comparable way to GPC-B-specific antibodies such as 37.7H; however, FP-1 preferentially engages the fusion peptide as its primary GPC contact whereas GPC-B antibodies typically have more extensive footprints and further engage the pocket formed by the C-terminal helix, HR1d helix, and the flexible loop upstream of β11 (Figures 4C and Figure S6B).^27,28^ When bound, FP-1 is flanked by N79, N99, and N373 glycans and is situated on the opposite side of the GP protomer relative to Base-1. While its peptidic interactions are localized to the GP2 subunit, the N79 glycan seemingly has a vast network of interactions with the LC FWR3 with contacts likely extending to the CDRL2 loop. These glycan interactions also serve as the only GP1 interactions with FP-1. The most notable putative contacts are between the CDRH3 and fusion peptide. Specifically, our model suggests residues 269–275 of the fusion peptide are interacting with FP-1’s CDRH2 and CDRH3. Additional engagement by the CDRL3 with residue K272 seems probable. Recent studies have shown multiple conformations of LASV’s fusion peptide are possible, with unbound (whether by matriglycan or antibodies) GPCs residing flexibly in the space between the HR1a helix, HR1d helix, and the HR2 loop.^30,31^ We note that binding by FP-1 seems to promote the fusion peptide conformation where residues 260-268 extend towards the interior trimeric interface, as has been observed when GPC is bound by other NAbs such as 25.10C and 37.7H. While we are yet unable to confirm, it is reasonable to suspect that FP-1 would have the potential to mature towards a neutralizing phenotype as has been reported with many GPC-B mAbs.

### Analysis of recombinant LASV GPC-induced antibody responses – the interior epitopes

Considering that GPCysR4-I53-50A trimers destabilize in the presence of pAbs, it is conceivable that the same would happen *in vivo*. This would result in the trimer exposing its internal and non-neutralizing epitopes on GP1 and GP2. As they represent a considerable peptidic surface on a highly glycosylated protein, these internal sites would likely induce a dominant antibody response. To investigate if these interior epitopes were being targeted, we performed EMPEM experiments using a pool of week 30 pAbs from rabbits 187-192 which were incubated with monomeric GP pre-complexed with 25.10C Fab (Figure 5A). This Fab was used as a fiducial marker to aid in determining the orientation and alignment of the stalky, featureless monomer. 2D classifications of extracted single particles revealed classes with two equatorial Fabs binding opposite sides of the monomer. In most classes, an additional Fab targeted the top or bottom of the GP monomer. Considering one of the two opposing equatorial Fabs includes 25.10C, the 2D classes suggest interior-targeting responses were elicited in these rabbits.

**Fig. 5.**
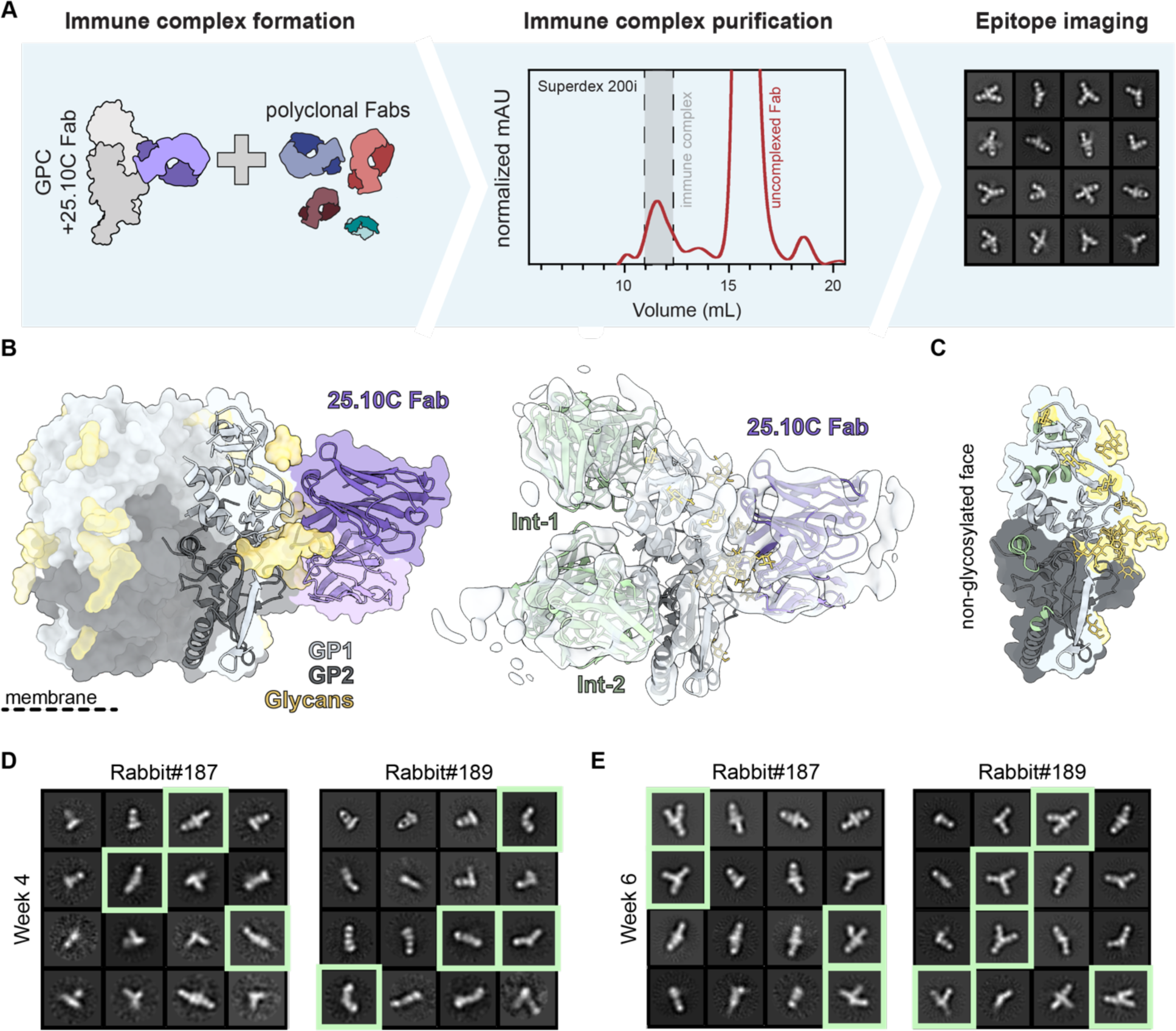
EMPEM experiments with GP monomers reveals responses targeting the trimer interior. **(A)** Schematic of monomeric GP EMPEM employing 25.10C Fab as a fiducial marker. **(B)** Monomeric GP cryo-EMPEM reveals the presence of interior binding pAbs Int-1 and Int-2. GP monomer with 25.10C Fab (PDB: 7TYV^40^) shown in trimeric context (left). Cryo-EM density map enabled docking of models of GPC monomer with 25.10C Fv (PDB: 7TYV^40^) and additional rabbit Fvs (PDB: 7SGF^30^). **(C)** Putative epitope footprints of Int-1 and Int-2 mapped onto GPC monomer show their preference for interior, non-glycosylated GPC surface. **(D and E)** 2D class averages of nsEMPEM experiments with monomeric GP pre-complexed with 25.10C Fab. Classes shown are the selected classes from two iterations of 2D classification that have been reclassified to 16 classes. **(D)** 2D class averages of monomeric GP nsEMPEM experiments with week 4 serum from rabbit 187 and 189. Classes with clear interior-binding pAbs are highlighted with green squares. **(E)** 2D class averages of monomeric GP nsEMPEM experiments with week 6 serum from rabbit 187 and 189. Classes with clear interior-binding pAbs are highlighted with green squares.

We used single-particle cryo-EM to provide more detail on these presumed interior-targeting pAbs (Table S3). While the overall resolution of our map was limited due to the small size of GP monomer and the sample’s vast heterogeneity, we were able to generate a map with adequate resolution to confidently dock in a GP monomer bound to 25.10C (PDB: 7TYV^40^). As hypothesized, we observed two interior binding pAbs, which we designated Int-1 and Int-2, targeting surfaces that would be inaccessible in the context of trimeric GPC (Figure 5B). Int-1 binds near the apex of the GP monomer and makes clear contacts with the α1 helix (aa 120–127) with contacts likely extending to α2 (aa 130–144). Int-2 is presumably the equatorial response we observed by nsEMPEM and resides opposite the 25.10C epitope. It appears to predominantly engage the HR1d helix (aa 347–358) but likely also contacts the loop just downstream of HR1d (aa 359– 362) and the beginning of the C-terminal helix HR2 (aa 398–402). Both pAbs show specificity for the non-glycosylated face of the exposed GP interior surface (Figure 5C). Interestingly, the interior-binding pAbs targeted similar epitopes as those putatively described for non-NAbs elicited by healthy LASV-seropositive human survivors from Sierra Leone and Nigeria.^15^ Int-1 shares its epitope with GP1-B Abs, which were originally thought to target residues between 119–134 while Int-2 engages with residues belonging to the presumptive GP2-B class of Abs.

Having confirmed the identity of these pAb responses by cryo-EM, we continued to perform nsEMPEM experiments with bleeds from earlier time points (Figures 5D and 5E). Complexes of GP monomer and 25.10C Fab were incubated with week 4 and 6 pFabs from rabbit 187 and 189 and imaged. At week 6, substantial Int-1 and Int-2 responses were observed in both rabbits, signified by the typical Y-shaped complexes in the 2D classes (Figure 5E). Even for the week 4 pFabs, for which we observed no responses to trimeric GPC, complexes of Int-2 pFabs and monomeric GP-25.10C Fab were discernible amongst the 2D classes (Figure 5D). Together these data suggest that interior-targeting responses dominate the immune response to GPCysR4-I53-50A trimers, showing up several immunizations before the elicitation of responses targeting neutralizing epitopes.

### Analysis of antibody responses to a GPC-derived VLP

Following our investigation of the pAb responses to rabbits vaccinated with recombinant GPC, we extended our EMPEM analysis to a potential vaccine candidate where GPC is embedded in the membrane. A previously described GPC-derived VLP, generated by Madin-Darby canine kidney (MDCK) cells stably expressing fullp-length GP and matrix protein, elicited remarkably strong autologous neutralization titers after four immunizations in a two-rabbit animal study (Figure 6A).^52^ Our efforts focused on rabbit 350, which demonstrated the highest neutralization titers of the two and exhibited prophylactic and therapeutic in vivo efficacy in lethal LASV infection model^52,64^. In a direct comparison with sera of GPCysR4-I53-50A-immunized rabbits, rabbit 350 induced an approximately 15-fold higher authentic neutralization titer than rabbit 187, the best neutralizer of the six rabbits (rabbit 187)(Figure S7A). In agreement with our findings that the base epitope is inaccessible on membrane-embedded GPC, we observed no base-targeting responses in nsEMPEM experiments with pAbs from rabbit 350 (Figure S7B). Rather, we observed what seemed to be several responses targeting the trimer apex. Furthermore, nsEMPEM with GP monomer revealed a pAb response bound to the exterior of GP (likely one of the apex-targeting pAbs) but no clear interior-targeting pAbs (Figure S7C).

**Fig. 6.**
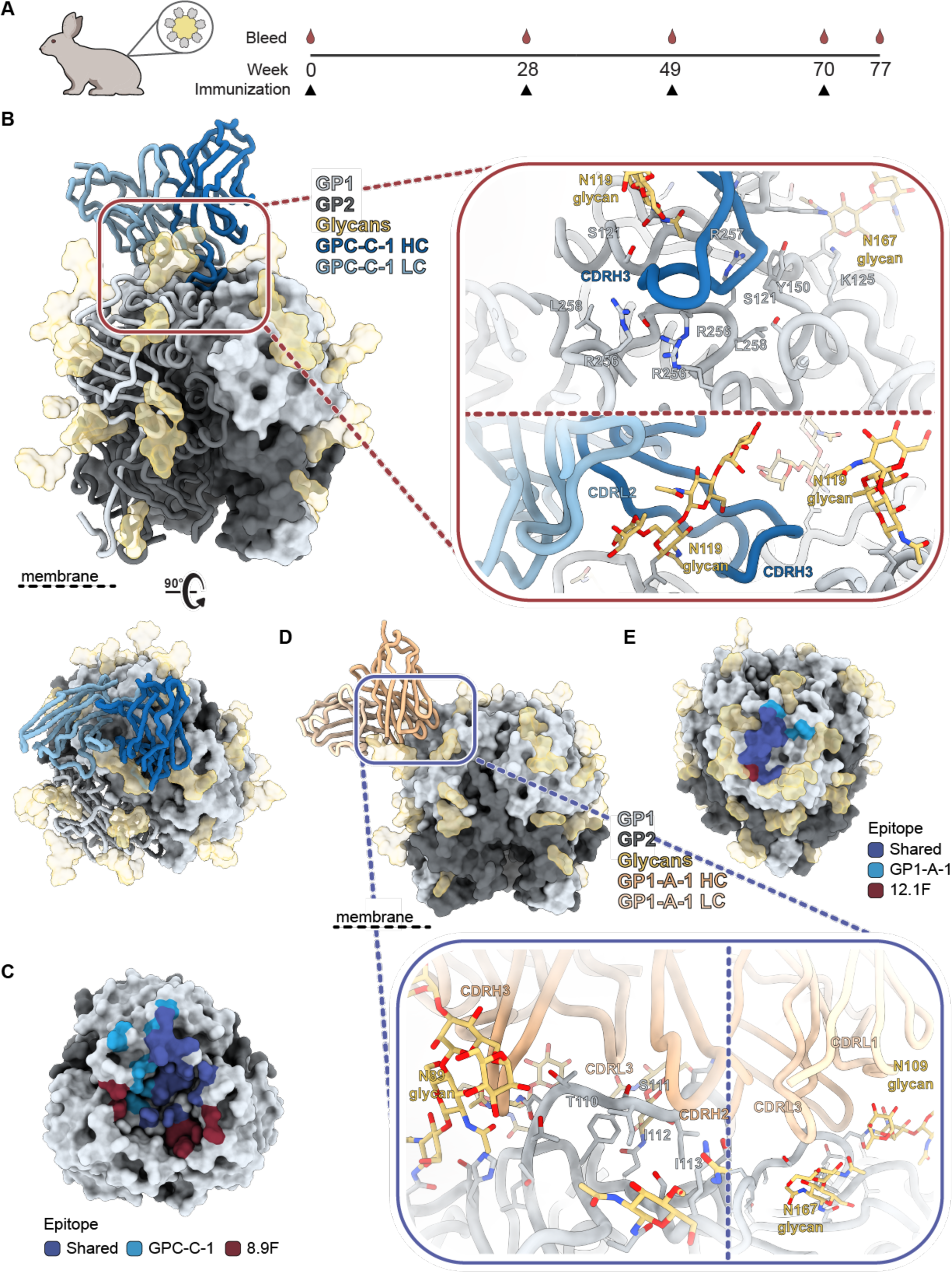
GPC-derived VLP immunization in rabbits induces antibody responses to the GPC-C and GP1-A epitope. **(A)** Immunization scheme of the rabbits immunized with a GPC-derived VLP, showing a prime at day 0, and boosts at day 28, 49 and 70, as recently described.^52^ **(B)** Atomic model of GPC-C-1 Fab bound to GPCysRRLL-I53-50A as determined by cryo-EM. Inset depicts putative interactions at the epitope-paratope interface which are highlighted and shown as sticks. **(C)** Epitope footprint comparison between GPC-C-1 and 8.9F Abs. Epitope-Analyzer^95^ was used to determine (putative) contact residues using a cut-off of 4 Å between epitope and paratope for 8.9F and 6 Å for GPC-C-1 (poly-alanine model). **(D)** Atomic model of GP1-A-1 Fab bound to GPCysRRLL-I53-50A as determined by cryo-EM. Inset depicts residues on GPC, shown as sticks, that likely make contacts with GP1-A-1. **(E)** Epitope footprint comparison between GP1-A-1 and 12.1F Abs. Epitope-Analyzer^95^ was used to determine (putative) contact residues using a cut-off of 4 Å between epitope and paratope for 12.1F and 6 Å for GP1-A-1 (poly-alanine model).

Given the absence of base-proximate responses, we performed cryo-EM using GPCysRRLL-I53-50A pre-complexed with 18.5C Fab. This led to the generation of two maps with a resolution of 2.8 Å each: one map highlighting a GPC-C response, the other a GP1-A response (Figures S3 and S4, Table S3). Upon closer inspection of the reconstructed model of the former, it became evident that the pFab, which we designate GPC-C-1, interacts with GPC in a manner strikingly similar to the bNAb 8.9F (Figure 6B and C). Firstly, GPC-C-1 employs an unusually extended CDRH3 (26 amino acids) to reach into the apical cavity of the trimer, establishing contacts with all three protomers. Secondly, interactions with the glycan at position N119 were observed. Lastly, like 8.9F, the light chain (LC) makes contacts with only one of the three protomers, also engaging the N119 glycan. Due to the high resolution of this map, we were able to speculate on the nature of certain GPC-pFab interactions where the pFab side chains were well-resolved (Figure S8A). We note, for instance, a likely aromatic density at position 100K in the CDRH3, making contacts with glycan N119. Additional aromatics can be observed at position 100L in the CDRH3, 32 in LC FR2, and and 93 in LC FR4 forming hydrophobic interactions with F117 in GPC. Furthermore, through what we speculate are arginine residues at positions 100A and 100D, the CDRH3 presumably forms cation-pi interactions with GPC’s Y150, a residue that has been shown to be critical for matriglycan binding. Based on our model, we suggest that GPC-C-1 played a substantial role in the observed neutralization in this rabbit by directly inhibiting the binding of GPC to matriglycan.

The generated model of the GP1-A-like response revealed that the pFab, designated GP1-A-1, is positioned in-between glycans N89, N109, 119, N167, and 224, and uses both its HC and LC to engage GP1. Besides several surrounding putative contacts with S91, D156, and K161, GP1-A-1 mainly targets the loop that extends over β5-β8 (aa 107-115) using its CDRH2, CDRH3, and CDRL3 (Figure 6D). Beyond these peptidic contacts, there seems to be extensive interactions between GP1-A-1 and glycans N89, and, to a lesser extent N109. We observed, for instance, putative aromatics in CDRH3 making contacts with glycan N89 as well as in CDRL3 and CDRL1 engaging with glycan N109 (Figure S8B). An interaction between glycan N167 and CDRL3 seems probable but likely only plays a minor role in the epitope-paratope interface. The way GP1-A-1 engages GPC is remarkably similar to the previously characterized NAb 12.1F and this is emphasized by the clear overlap in their epitope footprints (Figure 6E).^31^ 12.1F also primarily engages residues between amino acids 107 and 115 and makes contacts with glycans N89 and N109.^29,31^ As a result, 12.1F neutralizes LASV by inhibiting the conformational change that GP1 undergoes at late endosomal pH (5.5) and is required for LAMP-1 binding. Thus, unlike GPC-C-1, GP1-A-1 would need to withstand a low pH environment to confer neutralization.

## Discussion

The isolation of highly potent and protective NAbs from Lassa fever survivors demonstrates that the humoral immune response is capable of targeting sites of vulnerability on LASV’s highly glycosylated GPC, providing promise for the development of LASV vaccines aimed at inducing NAb responses. Nevertheless, induction of NAb responses by the various LASV vaccine efforts so far has been absent to poor, and their elicited Ab-response poorly understood. Detailed characterization of vaccine-induced antibody responses may be an important step in finding the bottlenecks for NAb development and designing more effective LASV vaccines. Serum ELISAs have been the standard to characterize vaccine-induced humoral responses, but they have generally been performed with non-stabilized, heterogeneous populations of GPC. Furthermore, as we show here, even with prefusion-stabilized GPC trimers, antibody-induced disassembly of GPC confounds interpretation. To address these shortcomings, we presented an EMPEM pipeline for LASV that enables highly detailed structural characterization of responses to both trimeric and monomeric forms of prefusion GPC. By testing our pipeline on sera from rabbit(s) previously vaccinated with GPC-I53-50A and a GPC-derived VLP, we identified several on- and off-target responses, including previously uncharacterized base and fusion peptide responses. These insights have implications for the development of next-generation recombinant GPC immunogens and may fuel the generation of vaccine candidates that have eliminated the exposure of undesired epitopes, paving the way for subdominant neutralizing responses to emerge.

EMPEM with sera from rabbits vaccinated with a recombinant GPC protein revealed that polyclonal antibody responses consistently target a previously undescribed epitope that is located at the C-terminal helix and N-terminal loop. This base epitope is part of a larger surface at the bottom of the trimer that is close to the membrane and, consequentially, lacks glycans. EMPEM and neutralization assays with monoclonal antibodies indicated that these base-targeting responses are non-neutralizing. We note a clear parallel to the base responses elicited by recombinant prefusion-stabilized HIV-1 Env trimers like the prototypic BG505 SOSIP.664. Over the years EMPEM studies have shown the consistency and immunodominance of base responses and have guided the redesign of Env trimers that attempt to shield this irrelevant neo-epitope.^53,57,60,65–68^ The inability of base-targeting mAbs to bind membrane-embedded GPC as well as the absence of base-targeting responses in the VLP-immunized rabbit suggest that the base epitope constitutes a neo-epitope generated by truncation of the transmembrane domain on recombinant GPC. This reveals a clear shortcoming of recombinant GPC vaccines. Still, future EMPEM studies on more VLP-immunized subjects, as well as those immunized with other vaccine modalities that use membrane-embedded GPC (i.e. mRNA, DNA or vector-based vaccines), will need to be performed to affirm if this neo-epitope is merely a disadvantage of recombinant GPC vaccines. Nevertheless, informed by our details on the epitope-paratope interface, this highly immunogenic off-target epitope should be ameliorated in the next iteration of recombinant GPC immunogens. For example, introduction of glycans in the C-terminal helix or N-terminal loop could enhance shielding of this epitope. Alternatively, as trimeric GPC ectodomains currently require stabilization domains, adjusting the rigidity and length of the C-terminal linkers to their trimerization scaffolds could occlude accessibility of the base.

Another off-target response we observed exploits the interior of the GPC trimer. As this epitope would be inaccessible in the trimeric conformation of prefusion GPCysR4-I53-50A, these responses indicate either that the GPC is disassembling *in vivo* or the GPC undergoes antibody-facilitated disassembly. This latter phenomenon has been observed extensively for influenza and HIV-1 where specific antibody responses have been shown to facilitate hemagglutinin and envelope protein dissolution, respectively.^69–71^ Our observations that interior-targeting responses were already elicited after the first immunization suggest that the GPCysR4-I53-50A readily disassemble *in vivo* before the emergence of other responses to the exterior of the trimer. As a result, GPC exposes its largest non-glycosylated surface. According to our analyses, this interior surface contains epitopes that are the dominant target during early immunizations and likely compete with B cells in germinal centers that recognize functional epitopes. Our monomer EMPEM data also correspond to previous reports of non-NAbs targeting the GP1-B and GP2-B epitopes.^15^ We offer structural definition to these antibody clusters with data that strongly suggests these epitopes are present only on disassembled, non-trimeric GPCs. The fact that these mAbs were isolated from Lassa fever survivors, suggests that natural infection presents GPC interiors, though the mechanism by which this happens is still unknown. Regardless, our results imply that fusing GPC to a trimerization domain is insufficient and further stabilization of the trimeric interface should dictate the design of the next generation of recombinant GPC vaccines.

Our EMPEM work also revealed vaccine-induced pAb responses remarkably similar to previously characterized NAbs isolated from Lassa fever survivors, supporting the use of rabbits as vaccination models. GPCysR4-I53-50A induced a response in two out of six rabbits that correlated strongly with neutralization and which we demonstrated parallels the canonical human GPC-A NAbs. With its high similarity to GPC-A-targeting NAbs 36.1F and 25.10C, GPC-A-1 likely neutralizes LASV by blocking LAMP-1.^40^ While the FP-1 pAb response did not fully mirror the defined GPC-B epitope, it engages with some of the same residues that are targeted by potent GPC-B NAbs.^27,28^ Focusing heavily on the fusion peptide, we speculate that the FP-1 response may have the potential to mature towards neutralization, perhaps by increasing its affinity or making it resilient to acidic pH. Interestingly, the GPC-derived VLP, which induced markedly more potent neutralizing responses than GPCysR4-I53-50A, elicited very different on-target responses. This vaccine induced responses very similar to the GPC-C and GP1-A NAbs, 8.9F and 12.1F, respectively; two of the three NAbs that make up Arevirumab-3.^29,43^ The engagement of GPC-C-1 with the native-cleavage site reinforces earlier suggestions that these residues are necessary for the elicitation of apex-targeting antibodies and provides one explanation for the differences in NAb responses between the vaccine modalities.^29^ Another explanation may be differences in epitope presentation (i.e. valency of immunogen display and accessibility of epitopes) between the two vaccines. It is tempting to perceive the remarkable neutralization titers this rabbit generated in the context of the VLP’s inability to induce off-target responses to the base and interior. It should be considered, however, that this immunogen required an extensive prime-boost regimen to achieve its potent neutralization. We do note that membrane-embedded GPC’s are generally poorly cleaved (the GPC-derived VLP is no exception) and may present immunogenic off-target responses that are not found on cleaved trimers. EMPEM experiments with uncleaved forms of GPC could shed light on the role these epitopes play in the low immunogenicity of vaccines that present membrane-embedded GPC.

In conclusion, we have presented direct high-resolution visualization of vaccine-induced humoral immune responses to prefusion GPC in its trimeric and monomeric conformational states. By identifying on- and off-target responses, we have highlighted sites of vulnerability on this densely glycosylated immunogen and revealed potential bottlenecks to the elicitation of NAb responses. Our demonstration of the immunodominance of the trimer base and interior not only has strong implications for vaccine design of recombinant GPC immunogens but also hints at the advantage of vaccines that present GPC on a membrane. As more attention is drawn to combatting Lassa fever and several vaccine candidates are moving to the clinic, the careful and robust assessment of humoral responses may be more important than ever. The work described here supports the use of our EMPEM pipeline as an integral component of serological assays which may considerably advance the understanding of induced antibody responses and, concomitantly, the development of a protective Lassa vaccine.

## Methods

### Construct design

The GPCysRRLL-I53-50A, GPCysR4-I53-50A, and GPCysR4 monomer constructs were generated as previously described and feature the strain Josiah GPC sequence (GenBank: NP_694870.1). Plasmids encoding TwinStrep-tagged or 6x-His-tagged 8.9F and 6x-His-tagged 18.5C were generated by introducing the respective tags in HC plasmids of 8.9F and 18.5C, as previously described.^30,31^

### Protein expression and purification

#### GPCysRRLL and GPCysR4 trimers and GPCysR4 monomer

GPCysRRLL-I53-50A, GPCysR4-I53-50A, and GPCysR4 monomers were transiently expressed in HEK 293F cells at a density of 1.0 × 10^6^ cells/mL. Ratios of 1:3 DNA to PEImax were used to facilitate co-transfection of plasmids encoding GPCysRRLL-I53-50A, GPCysR4-I53-50A, or GPCysR4 with plasmids encoding S1P or furin. A ratio of 2:1 GPC to protease plasmid DNA was used. HEK 293F cells were cultured and kept shaking at 125 rpm for five days at 37°C and 8% CO2. Cultures were harvested after six days and GPC-I53-50As and GPCysR4 monomers were purified via StrepTag purification using gravity columns and StrepTactin 4Flow resin (IBA Life Sciences), following the manufacturer’s protocol. The proteins were eluted using 1X BXT buffer (IBA Life Sciences) and buffer exchanged to 1X TBS using 100 kDa MWCO concentrators for trimers and 10 kDa MWCO concentrators for monomers (Millipore). As a final purification step, size exclusion chromatography was performed using Superdex 200 increase 10/300 GL columns (Sigma-Aldrich) with TBS as the running buffer. Fractions associated with the elution volume corresponding to trimeric or monomeric GPC were collected and concentrated using either 100 kDa or 10 kDa MWCO concentrator, respectively (Millipore).

#### 8.9F and 18.5C Fab

8.9F and 18.5C Fabs were expressed in HEK 293F cells using a transfection ratio of 2:1 LC to HC and a 3:1 PEI to DNA ratio. 8.9F HCs featured either a 6x-His tag or a TwinStrep tag while 18.5C HC encodes a 6x-His tag. Cells were cultured for six days as mentioned above before being harvested by centrifugation. Depending on the tag, Fabs were purified using StrepTactin 4Flow resin (IBA Life Sciences) or Ni-NTA resin (Thermo Scientific). TwinStrep tagged 8.9F was purified via gravity column before being eluted by 1X BXT (IBA Life Sciences) while 6x-His tagged 8.9F and 18.5C culture supernatants were rolled overnight with Ni-NTA beads at 4°C before using gravity columns to capture the resin. This resin was washed with 20 mM imidazole, 50 mM NaCl, pH 7.0 buffer before being eluted with a 500 mM imidazole, 50 mM NaCl, pH 7.0 buffer. All Fabs were buffer exchanged to TBS using 10 kDa MWCO concentrators (Millipore).

#### IgG

8.9F, 12.1F, 25.10C, and 37.7H IgGs were expressed as previously described.^30,31^ In brief, IgG plasmids were transfected at a HC to LC ratio of 1:1 in HEK 293F cells. Cells were cultured and harvested by centrifugation five days post-transfection. The IgGs were purified by adding Protein G (Cytiva) or CaptureSelect IgG-Fc resin (Thermo Scientific) to the supernatant and rolling overnight at 4°C. Resin was captured by gravity column and IgG eluted with 0.1 M glycine at pH 2.0 into 1 M Tris pH 8.0. All IgGs were immediately buffer exchanged to TBS using 30 kDa MWCO concentrators (Millipore).

#### GPCysRRLL-I53-50A+8.9F Fab complexes

Complexes of 8.9F and GPCysRRLL-I53-50A were expressed and purified directly to remove the need for final size exclusion chromatography steps. In these cases, 500 µg purified 8.9F Fabs with a TwinStrep tag were added to transfected HEK 293F cells (see above) expressing GPCysRRLL-I53-50A with an Avi-His tag at day three post-transfection. After harvesting cells at day six post-transfection, 8.9F-GPCysRRLL-I53-50A complexes were purified using StrepTactin 4Flow resin as mentioned previously and then buffer exchanged to TBS using 100 kDa MWCO concentrators (Millipore).

Complexes of 8.9F and GPCysRRLL-I53-50A were also generated using parallel HEK 293F transfections. In these cases, HEK 293F cells were split in a 1:4 ratio and grown to 1.0 × 10^6^ cells/mL. The smaller batch of cells were transfected with a 2:1 ratio of 8.9F LC and HC plasmid DNA. The HC plasmid DNA featured a TwinStrep tag. The larger batch of cells was transfected with GPCysRRLL-I53-50A with an Avi-His tag as previously described. On day three post-transfection, the HEK 293F cells expressing 8.9F Fab were added to the culture expressing GPCysRRLL. Cultures were harvested at day six post-transfection and complexes were purified using StrepTactin 4Flow resin (Thermo Scientific) and eluted with 1X BXT. Complexes were immediately buffer exchanged to TBS using 100 kDa MWCO concentrators (Millipore).

#### GPCysRRLL-I53-50A+18.5C Fab complexes

Complexes of 18.5C and GPCysRRLL-I53-50A were made by mixing 50 µg of purified 18.5C Fab and 20 µg of purified GPCysRRLL/R4-I53-50A followed by an incubation at 4°C for 1 hour.

### Isolation of LAVA05 and LAVA06

Rabbit B cells were isolated by flow cytometry (BD, FACSAria) using mouse-anti-rabbit IgG-PE (Southern Biotech) on frozen peripheral blood mononuclear cells (PBMC) samples from animals that were vaccinated with the recombinant GPCysR4-I53-50A.^30^ To exclude I53-50A-binding B cells we included a SARS-CoV-1 Spike probe containing I53-50A-biotin which was labeled with streptavidin AF647 and streptavidin BV421 (both Biolegend). After a 2 day culture with interleukin (IL)-21 and irradiated mouse fibroblast (L cells) expressing CD40-ligand in DMEM F12 media (Gibco), the B cells were immortalized by retroviral transduction with BcL6, Bcl-XL and the marker gene GFP, as previously described.^72,73^ Subsequently, the cells were cultured at 1 cell per well in 96 well plates. After 10 days culture supernatant containing secreted monoclonal antibody were screened for binding to GPCysR4-I53-50A (strain Josiah) using a custom Luminex assay similar to previously described.^74^ In short, viral antigens were covalently coupled to Luminex Magplex beads with a two-step carbodiimide reaction at a ratio of 75 µg protein to 12.5 million beads. Beads and B cell culture supernatant were incubated overnight, followed by detection with mouse-anti-rabbit IgG-PE (Southern Biotech). Read-out was performed on a Magpix (Luminex). Positive clones were collected and a cell pellet was stored for VH and VL sequence determination and cloning.

### Antibody cloning

First, a combined reverse transcription (RT) and first PCR reaction was performed on 1 µl of a 1 in 100 diluted freeze-thawed cell pellet using the SCRIPT HF RT-PCR Kit (Jena Bioscience), according to the manufacturer’s protocol. For the first PCR we included 0.5 µl of primers from a 10 mM stock as described in Table S1 of McCoy et al.^75^ The RT reaction was performed for 1 hour at 60°C and 5 min at 5°C after which the first PCR reaction was started immediately for 40 cycles at 95°C for 10 s, 20 s at 55°C and 1 min at 72°C and ending with 2 min at 72°C. Subsequently a second PCR was performed which included primers with Gibson overhang in the leader sequence and start of the hinge region. These Gibson sequences are necessary for cloning of the VH and VL into our rabbit IgG1 expression vectors which are based on the pFUSE-rIgG-Fc (InvivoGen) vectors.^75,76^ For this second PCR, 0.4 µL of the first PCR was used in a reaction using HotStarTaq polymerase (Qiagen) for 5 min at 95°Cand 30 cycles of [30 s at 94°C, 30 s at 60°C, and 1 min at 72°C], and 10 min at 72°C. Gibson cloning was then used to integrate the amplified heavy and light chain V(D)J variable regions in mammalian cell expression vectors containing the rabbit constant regions. This was done by mixing 1 µL of expression vector, 1 µL of the second PCR product, and 2 µL of home-made Gibson mix (T5 exonuclease (0.2U; New England Biolabs), Phusion polymerase (12.5U; New England Biolabs), TaqDNA ligase (2000U; New England Biolabs), Gibson reaction buffer (0.5 grams PEG-8000; Sigma Life Sciences), 1 M Tris/ HCl pH 7.5, 1 M MgCl2, 1 M DTT, 100 mM dNTPs, 50 mM NAD + (New England Biolabs) MQ) and incubating the mix for 60 min at 50°C.

### Antibody binding to membrane expressed GPC

One day prior to transfection with PEIMax of 8 µg of full length native GPC (strain Josiah) expressed in the pPPI4 vector, 3×10^6^ HEK293 cells in 8 mL DMEM were seeded per petridish. Cells were harvested after approximately 36 hours and frozen at 2×10^6^ per vial. For the flow cytometry assay (FACS Symphony, BD) non-transfected HEK293 cells were labeled with the CFSE CellTrace and the Soromba expressing cells were labeled with Violet CellTrace (Invitrogen). After washing, the cells were pooled including non-stained HEK293 cells expressing native GPC (strain Josiah). Next the cells were incubated with a dilution range starting at 10 µg/mL of purified monoclonal antibodies either from human or rabbit origin. Antibody binding was detected either with mouse-anti-human or rabbit IgG-PE (Invitrogen) and detected using the FACS Symphony (BD). Analysis was performed using FlowJo software.

### Antibody binding by biolayer interferometry

Ab binding to recombinant GPCyRRLL-I53-50A was determined using an Octet Red96 instrument (ForteBio). 8.9F, 12.1F, 25.10C, and 37.7H IgG were diluted to concentrations of 10 µg/mL and loaded onto Protein A sensors (Sartorius) to a binding signal of 1.0 nM. Sensors were dipped into wells containing 120 nM GPCysRRLL-I53-50A for 600 s then dipped into kinetics buffer to measure dissociation for 600 s. To measure binding by base-specific mAbs, his-tagged 8.9F Fab at 10 µg/mL were loaded onto NiNTA sensors, after which sensors were dipped into wells containing 120 nM of GPCysRRLL-I53-50A for 600s. After a dissociation step of 600s (only a minor off-rate was observed witch reached a plateau within 600s), the sensor was dipped into a well containing 10 µg/mL LAVA05 or LAVA06 for 600s and then dipped into kinetics buffer to measure dissociation for 300 s. All assays were performed at 25°C with dilutions made in the manufacturer-recommended running buffer (PBS, 0.1% BSA, 0.02% Tween20, pH 7.4). Data was analyzed using the Octet Data Analysis software where kinetics buffer references were subtracted from all data.

### Differential scanning fluorimetry

Thermostability of GPCysR4-I53-50A, GPCysRRLL-I53-50A, and GPCysRRLL-I53-50A in complex with 8.9F was determined with a nano-DSF NT.48 (Prometheus). GPC proteins or complexes were diluted to 0.5 mg/mL and loaded into high sensitivity capillaries. The assay was run with a linear scan rate of 1°C/min and 80%-100% excitation power. The first derivative of the ratio of tryptophan fluorescence wavelength emissions at 350 and 330 nM were analyzed to determine thermal onset (*T_onset_*) and denaturation (*T_m_*) temperatures using the Prometheus NT software.

### Fab preparation for EMPEM studies

First, IgGs were isolated from sera using 2 mL CaptureSelect IgG-Fc resin slurry (Thermo Scientific) per 1 mL rabbit sera. Sera and resin were gently mixed at 4°C for a minimum of two overnights. After, resin was collected by a gravity-flow column and washed three times with 1X PBS, pH 7.4 to remove IgG-depleted sera. IgGs were eluted from resin twice with 9 mL of 0.1 M glycine, pH 2.0 buffer and immediately neutralized with 1mL 1 M Tris, pH 8.0. The elutions were then buffer exchanged to PBS using a 30 kDa MWCO concentrator (Millipore). IgGs were digested to Fab following previously described protocols (Han et al., 2019). In summary, IgGs were digested by incubating the IgGs with immobilized papain (Thermo Fisher) in freshly prepared digestion buffer (20 mM sodium phosphate, 10 mM EDTA, 20 mM L-cysteine, pH 7.4). IgGs were digested at 37°C for 4-5 hours before the reaction was quenched using iodoacetamide at a final concentration of 0.03 M. 2 mL of CaptureSelect resin slurry was added to digested IgG samples and incubated for a minimum of 2 hours at 4°C. CaptureSelect resin bound to Fc were separated from Fab mixtures by a gravity-flow column. The Fc-depleted flow-through was buffer exchanged to TBS using 10 kDa Amicon MWCO concentrators. For sera from rabbits immunized with GPCysR4-I53-50A, to remove I53-50A-specific antibody responses, the polyclonal Fabs were incubated with BG505 SOSIP scaffolded on I53-50 (BG505-I53-50A; construct design and expression described previously^77^) at a ratio of 1 mg purified Fab to 400 µg BG505-I53-50A. The mixture was incubated overnight at room temperature and run through a 100 kDa MWCO concentrators (Millipore) so the flow-through was depleted of Fabs specific to the I53-50A scaffold. The flow-through was finally concentrated using a 10 kDa MWCO Amicon concentrator (Millipore).

### Preparation of LASV glycoprotein immune complexes with polyclonal rabbit Fabs

For nsEMPEM as well as cryo-EM with graphene oxide grids, immune complexes were generated by mixing 20 μg of GPCysRRLL-I53-50A pre-complexed with 8.9F or 18.5C Fabs, as described above, with 600 μg of purified polyclonal Fab (in the case of GPCysR4-I53-50A sera, already depleted of I53-50A-specific Fabs). The mixture was incubated for 4 hours at 4°C. Immune complexes were then purified from unbound/excess Fab by size exclusion chromatography using a Superose 200 increase column with TBS as the running buffer. Fractions corresponding to immune complexes (∼9.8 mL–11.2 mL) were pooled and concentrated using a 100,000 MWCO centrifugal filter (Merck Millipore). To generate immune complexes for cryo-EM where UltrAuFoil R1.2/1.3 grids were used and consequentially a higher concentration of immune complex was required. For these experiments we complexed 200 μg of GPCysRRLL-I53-50A, pre-complexed with 8.9F Fabs, with 2000 μg of purified polyclonal Fab (already depleted of I53-50A-specific Fabs).

### Negative stain electron microscopy

Carbon-coated Cu-grids (400-mesh) were glow-discharged for 25 s at 15 mA using a PELCO easiGlow instrument (Ted Pella, Inc.). Samples were diluted to 15 μg/mL in TBS and loaded onto the grid for 30 s. After blotting off the sample, the grid was immediately stained with 2% (w/v) uranyl formate for 20-40 s. After blotting off excess stain, grids were dried for at least 5 min. Imaging was performed on a Tecnai Spirit electron microscope (operating at 120 keV, nominal magnification was 52,000 X, resulting pixel size at the specimen plane was 2.05 Å). Electron dose was set to 25 e-/Å2 and the nominal defocus for imaging was −1.50 μm. The Leginon automated imaging software^78^ was used for data acquisition.

### Negative stain electron microscopy data processing

Particles were picked using Appion,^79^ after which Relion 3.0 was used for particles extraction, 2D/3D classification and 3D refinement. Extracted particles were subjected to one to two rounds of 2D classification to select for GPCysRRLL-I53-50A-8.9F Fab or GPCysRRLL/GPCysR4-I53-50A-18.5C Fab complexes with a pFab bound. As none of the GPCysR4-I53-50A-vaccinated rabbits showed any pFab binding to GPCysR4-I53-50A-18.5C complexes we did not continue to generate 3D maps from these 2D classes. In addition, as we directly pursued cryo-EM, we also did not generate 3D maps of the immune complexes from sera of rabbit 350. The following therefore only applies to GPCysRRLL-I53-50A-8.9F Fab complexed with pFab. Once a minimum of 30,000 particles were selected from 2D classification, 3D classification was performed using a GPCysRRLL-I53-50A-8.9F complex as an initial model. This model was generated by low-pass filtering the following ensemble: a GPCysRRLL-8.9F complex (PDB: 8ER3) to which we docked in a full Fab (PDB: 4ZTP^63^) so that the complex would contain the constant domain) and a correctly positioned I53-50A scaffold (PDB: 7SGE^30^). After a first round of 3D classification, output maps were aligned to the initial model in UCSF Chimera using the Volume Fit tool. Correct alignment of the, at this resolution, almost featureless GPC is aided by the asymmetrical binding of 8.9F to GPC and the lobe-like structures of the protomers. Particles of classes that showed pFab(s) bound to the same epitope clusters were combined and subjected to another round of 3D classifications. This process was repeated until a well-defined GPC complexed with 8.9F and a pFab was visible. The combined particle stack was then further processed using 3D auto-refinement. Composite models of GPC with pAbs bound, as seen in Figures 2 and 3, were generated in UCSF Chimera by fitting both the refined 3D map as well as a low-pass filtered smoothened map of GPC to the initial model. All density apart from the pFab was then erased from the refined 3D map.

### Authentic neutralization assay

Authentic LASV (strain Josiah, lineage IV) neutralization assays were conducted in the BSL-4 laboratory at the Institute of Virology, Philipps University Marburg, Germany. Rabbit sera were inactivated with complement for 30 min at 56°C and subjected to a two-fold dilution series, starting at a dilution of 16. The diluted sera were then incubated with 100 TCID50 virus for 60 min at 37°C. Subsequently, Vero E6 cell (ATCC) suspension was added, and the plates were incubated at 37°C with 5% CO2. Cytopathic effects (CPE) were assessed seven days post-infection, and neutralization titers were determined as the geometric mean titer (GMT) of four replicates.

### Cryo-EM grid preparation and data collection

UltrAuFoil R1.2/1.3 grids (Au, 300-mesh; Quantifoil Micro Tools GmbH) were plasma discharged using a PELCO easiGlow (Ted Pella Inc.) for 25 s at 15 mA. Fluoro-octyl maltoside or lauryl maltose neopentyl glycol (Anatrace) were added to protein samples immediately before their applications to grids at final concentrations of 0.02 w/v% and 5 µM, respectively. Grids were plunge-frozen using a Vitrobot Mark IV (Thermo Fisher Scientific) with chamber temperature set to 4°C and humidity to 100%. Blot times between 3–5 s, blot force of 1 and a wait time of 10 s were used for all samples. Sample concentrations were between 0.6–4.0 mg/mL. For cryo-EM of GPCysRRLL-I53-50A-18.5C complexes with pFab from the GPC-derived VLP vaccinee, Quantifoil R1.2/1.3 grids (Cu, 400-mesh; Electron Microscopy Sciences) coated with graphene oxide were used. Sample, at a concentration of 0.1 mg/mL was loaded directly on the grid (not plasma discharged) in the absence of detergent and plunge-frozen using a Vitrobot Mark IV (Thermo Fisher Scientific) with chamber temperature set to 4°C and humidity to 100%. Blot time of 2.5 s, blot force of 0 and a wait time of 30 s were used.

Grids were loaded into a 300 kV FEI Titan Krios or 200 kV Thermo Fischer Scientific Glacios (II). The Titan Krios was equipped with a K2 Summit direct electron detector camera (Gatan) while the Glacios (II) featured a Falcon 4 detector (Thermo Fisher Scientific). Data collection was automated using Leginon and EPU software, respectively.^80^ Additional information is noted in Table S3.

### Cryo-EM data processing

Image preprocessing was performed either with the Appion software package^79^ or CryoSPARC Live software.^78^ For the former, micrograph movie frames were aligned and dose-weighted with UCSF MotionCor 2 software^81^ before being transferred to CryoSPARC 3.2.^78^ When CryoSPARC Live software was used, image motion corrections and initial CTF estimations were performed directly within that software package before transfer to cryoSPARC 3.2 for further processing. Particles were picked and classified into 2D classes. Good initial 2D classes were chosen for template picking of particles. This particle stack was iteratively 2D classified to remove bad particle picks.

#### GPCysRRLL-I53-50A bound to 8.9F

We used an initial model generated in UCSF ChimeraX^82^ from low-pass filtered GPC structure PDB: 7SGD.^30^ Particles were initially refined using homogeneous and heterogenous refinements to select for 3D classes with a good distribution of orientations. After initial alignments, we performed a focused 3D classification of the 8.9F antibody and selected particles based on their resolution at the epitope-paratope interface. Iterative rounds of global and local CTF refinements paired with local refinements were used to obtain a high resolution map.^83^ All refinements were performed with C1 symmetry due to the asymmetric engagement of 8.9F Fab.

#### GPCysRRLL-I53-50A bound to pAbs from GPCysR4-I53-50A-vaccinated rabbits

A low pass filtered structure of the 8.9F-GPCysRRLL-I53-50A complex was used as the initial model. Particles were first aligned to their shared GPC features through homogeneous refinements. Aligned particles were symmetry expanded along the C3 axis in accordance with the trimeric nature of GPC and to simplify further processing. As such, further alignments were restrained by using only local refinement and 3D Variability analyses to prevent symmetry-related particle copies from aligning to themselves.^84^ 3D variability analyses employing sphere masks around putative epitope clusters were based on nsEMPEM data. Clusters with Fab densities were further refined using iterative local refinements paired with 3D variability analyses to obtain particle stacks with the best epitope-paratope density.

#### GPCysRRLL-I53-50A bound to pAbs from GPC-derived VLP-vaccinated rabbits

Heterogeneous refinement was first performed to classify and refine a homogeneous stack of particles, using an initial model generated from ab initio refinement of the 2D classified particle stack. The particle stack corresponding to a GPCysRRLL-I53-50A-18.5C complex with three pFabs bound (2x GP1-A-, 1x GPC-C-like response) was then subjected to a round of NU-refinement after which 3D classification was performed using a sphere mask around the epitope-paratope interface. This was done separately for both the GP1-A-like epitope-paratope interface as well as the GPC-C-like epitope-paratope interface. 3D classes with the highest resolution particles were selected and subjected to multiple rounds of NU-refinement, separately for both pFab responses, using a soft solvent mask around the GPCysRRLL in complex with the pFab in question.

#### GPCysR4 monomer bound to pAs

After initial 2D classification, we used Topaz particle picking to identify GP particles bound to pAbs.^85^ We used initial models generated from ab initio refinements of the refined particle stack. Heterogeneous refinements enabled us to determine a class of immune complexes bound by three Fabs, which we further refined using iterative NU-refine and local refine jobs.^86^ For the final refinement, we employed an initial model and mask that contained only low-pass filtered GP protomer with Fv regions of bound Fabs based on PDB: 7TYV. ^40^

### Atomic model refinement

Sharpened maps were used to build all final atomic models. All initial GPC models (derived from PDB: 7SGD^30^, PDB: 8EJD^30,31^, or 8ER3) were manually fit using Coot.^87,88^ The 8.9F initial model was generated using ABodyBuilder.^89^ The initial models for pAbs were derived from PDB: 7RA7. The model 7RA7 was manually fit into densities for pAbs and based on the experimental density, residues were added or deleted from CDR loops. Conserved residues were maintained in the initial pAb models for early rounds of Rosetta refinement^90^ followed by manual corrections in Coot using additional models (PDB: 4ZTP,^63^ PDB: 4ZTO,^63^ and PDB: 7SGF^30^) to guide placement of loops. Once the fit of pAbs with conserved side chains was feasible, we re-defined all amino acids in the antibody sequence to alanine. Model fit to map was validated using Molprobity and EMRinger in the Phenix software package.^91–93^ Orientations of all glycans was assessed and corrected using Privateer.^94^ The epitope-paratope interactions between GPCysRRLL and 8.9F/pAbs were determined using Epitope-Analyzer.^95^ A cut-off of 4 Å was used for the peptidic contacts between GPCysRRLL and 8.9F, whereas for the pAbs (poly-A models) a cutoff of 6 Å was used. Glycans were defined to be involved at the epitope-paratope interaction of 8.9F and GPCysRRLL if they were in close proximity (<4 Å) using UCSF ChimeraX’s contacts tool.^82^ Cα RMSD values comparing our 8.9F-GPCysRRLL-I53-50A structure to PDB: 7UOT was performed using UCSF ChimeraX’s MatchMaker tool.^82^ Values denoted in the text represent the pruned and unpruned pairs of all GP1 protomers with the 8.9F HC and LC and exclude glycans. Final atomic models were submitted to the Protein Data Bank with accession codes found in Table S3. Figures including atomic models were all produced using UCSF ChimeraX.^82^

## Acknowledgments

The authors thank Bill Anderson, William Lessin and Hannah Turner from The Scripps Research Institute for their help with EM experiments, and Gabriel Ozorowski for help with cryo-EM data processing. We thank Gotthard Ludwig and Sebastian Schmidt from the biosafety level 4 facility at the Philipps-University of Marburg for technical support. We thank Robin Shattock for kindly sharing the full-length native Josiah GPC plasmid. H.R.P. is supported by a David C. Fairchild Endowed Fellowship, Achievement Rewards for College Scientists Foundation, and NIH F31 Ruth L. Kirschstein Predoctoral Award 1F31Al172358. P.J.M.B. is supported by a Rubicon fellowship from the Netherlands Organisation for Scientific Research (NWO). Further support from the Vici fellowship from the Netherlands Organisation for Scientific Research (NWO; to R.W.S.); by the Fondation Dormeur, Vaduz (to R.W.S. and M.J.v.G.); by the Deutsche Forschungsgemeinschaft (DFG, German Research Foundation)-Projektnummer 197785619/SFB1021 (to T.S.); by NIH grant R01 AI171438 (to A.B.W.); and by the Bill and Melinda Gates Foundation through grant OPP1170236 (to A.B.W.) enabled this work.

## Author contributions

Conceptualization, P.J.M.B, H.R.P., and A.B.W.; methodology, P.J.M.B., H.R.P., T.B., and H.M.-K.; formal analysis, P.J.M.B., H.R.P., T.B.; investigation, P.J.M.B., H.R.P., T.B., H.N., S.K., J.A.B., W.L., and H.M.-K.; resources, R.W.S., T.S., M.J.v.G., and A.B.W.; writing – original draft, P.J.M.B., H.R.P., and A.B.W.; reviewing, editing, and other feedback,; P.J.M.B., H.R.P., T.B., J.A.B., R.W.S., T.S., M.J.v.G., and A.B.W, visualization, P.J.M.B, and H.R.P; supervision, T.B., R.W.S, M.J.v.G, and A.B.W.; project administration, P.J.M.B., H.R.P., and A.B.W.; funding acquisition, R.W.S., T.S., M.J.v.G., and A.B.W.

## Data and code availability

- Maps generated from the cryo-EM data are deposited in the Electron Microscopy Databank (http://www.emdatabank.org/) under accession IDs: EMD-28549, EMD-41712, EMD-41713, EMD-41714, EMD-41715, EMD-41716, EMD-43141, and EMD-43168. Maps generated from the nsEM data are deposited in the Electron Microscopy Databank (http://www.emdatabank.org/) under accession IDs: EMD-43174, EMD-43175, EMD-43176, EMD-43177, EMD-43178, EMD-43179, EMD-43180, EMD-43181, EMD-43182, and EMD-43183. Atomic models corresponding to these maps have been deposited in the Protein DataBank (http://www.rcsb.org/) under accession IDs: 8ER3, 8TYB, 8TYC, 8TYE, 8VCV, and 8VE8. The raw data reported in this study will be shared by the corresponding author upon request.
- This paper does not report original code.
- Any additional information required to reanalyze the data reported in this work paper is available from the lead contact upon request.

