## Supplemental Material for "Defining bottlenecks and opportunities for Lassa virus neutralization by structural profiling of vaccine-induced polyclonal antibody responses"

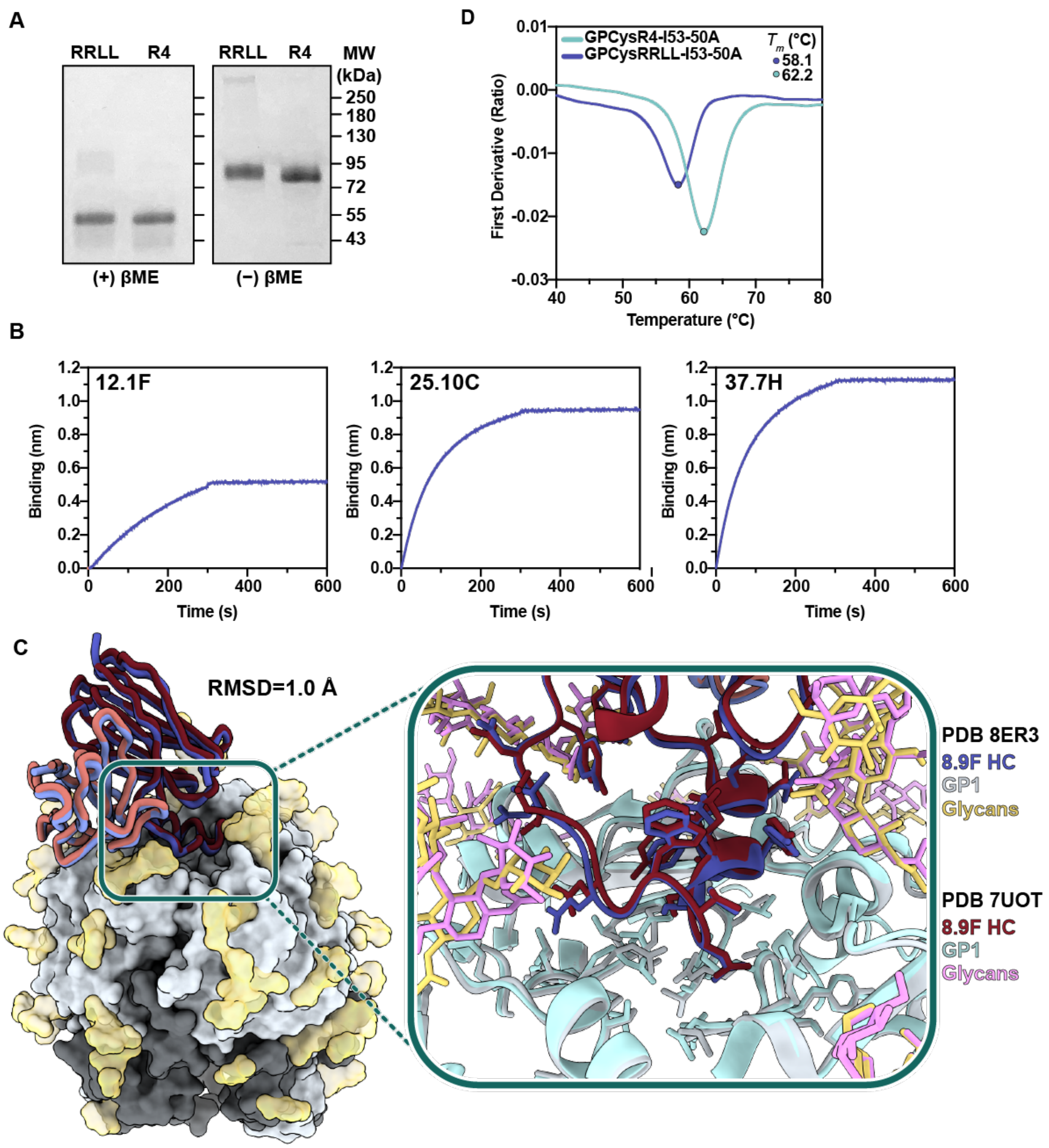


**Fig. S1. Characterization and comparison of GPCysRRLL-I53-50A, related to figure 1. (A)** SDS-PAGE comparison of cleavage between GPCysRRLL-I53-50A co-transfected with S1P and GPCysR4-I53-50A co-transfected with furin in reducing (left) and non-reducing (right) conditions. **(B)** BLI sensorgrams indicating immobilized 12.1F, 25.10C, and 37.7H IgGs binding to GPCysRRLL-I53-50A at concentration of 120 nM. **(C)** Comparison of GPCysRRLL-I53-50A bound to 8.9F with fully native GPC bound to 8.9F and 37.2D.[1] **(D)** Thermostability of GPCysRRLL-I53-50A and GPCysR4-I53-50A assessed by nanoDSF. Melting temperatures ($T_m$) are calculated as the inflection point (circles) of the ratio of signal at 350 and 330 nM. Each melting curve is a representative of triplicate curves with $T_m$ within ±0.1°C.

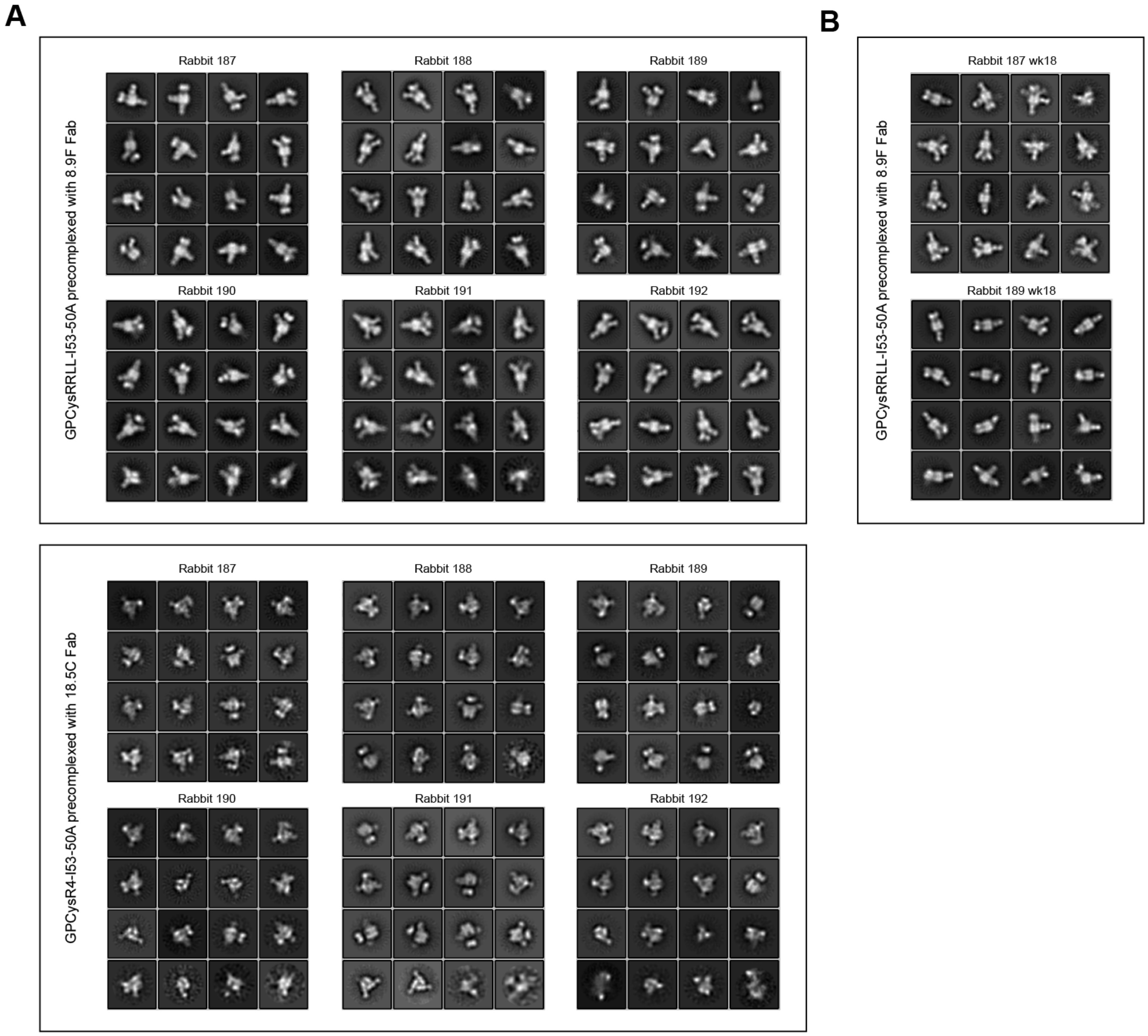


**Figure S2. 2D classes of selected particles for the generation of 3D maps, related to figure 2. (A)** 2D class averages of nsEMPEM experiments with week 30 pAbs from rabbit 187-192 using GPCysRRLL-I53-50A pre-complexed with 8.9F (top) and GPCysR4-I53-50A pre-complexed with 18.5C (bottom). Classes shown are the selected classes that were selected for 3D classification, reclassified to 16 classes. **(B)** 2D class averages of nsEMPEM experiments with week 18 pAbs from rabbit 187 and 189 using GPCysRRLL-I53-50A pre-complexed with 8.9F. Classes shown are the selected classes that were selected for 3D classification, reclassified to 16 classes.

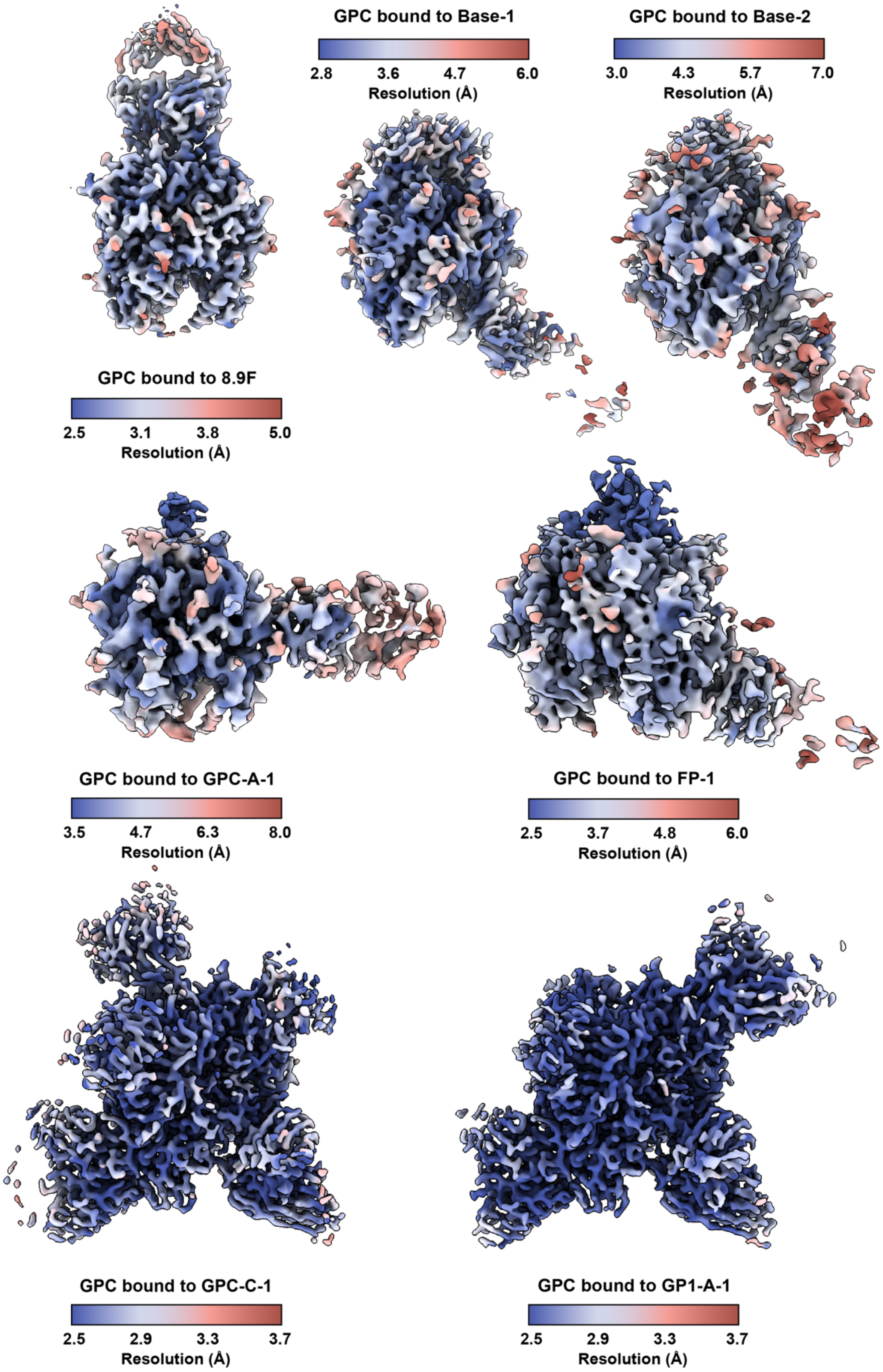


**Fig. S3: Local resolution plots for EM maps of LASV GPC and 8.9F, Base-1, Base-2, GPC-A-1, FP-1, GPC-C-1, and GP1-A-1, related to figures 3, 4, and 6**. Local resolutions calculated using cryoSPARC 3.2[2] using an FSC threshold of 0.143. Maps visualized in ChimeraX.[3]

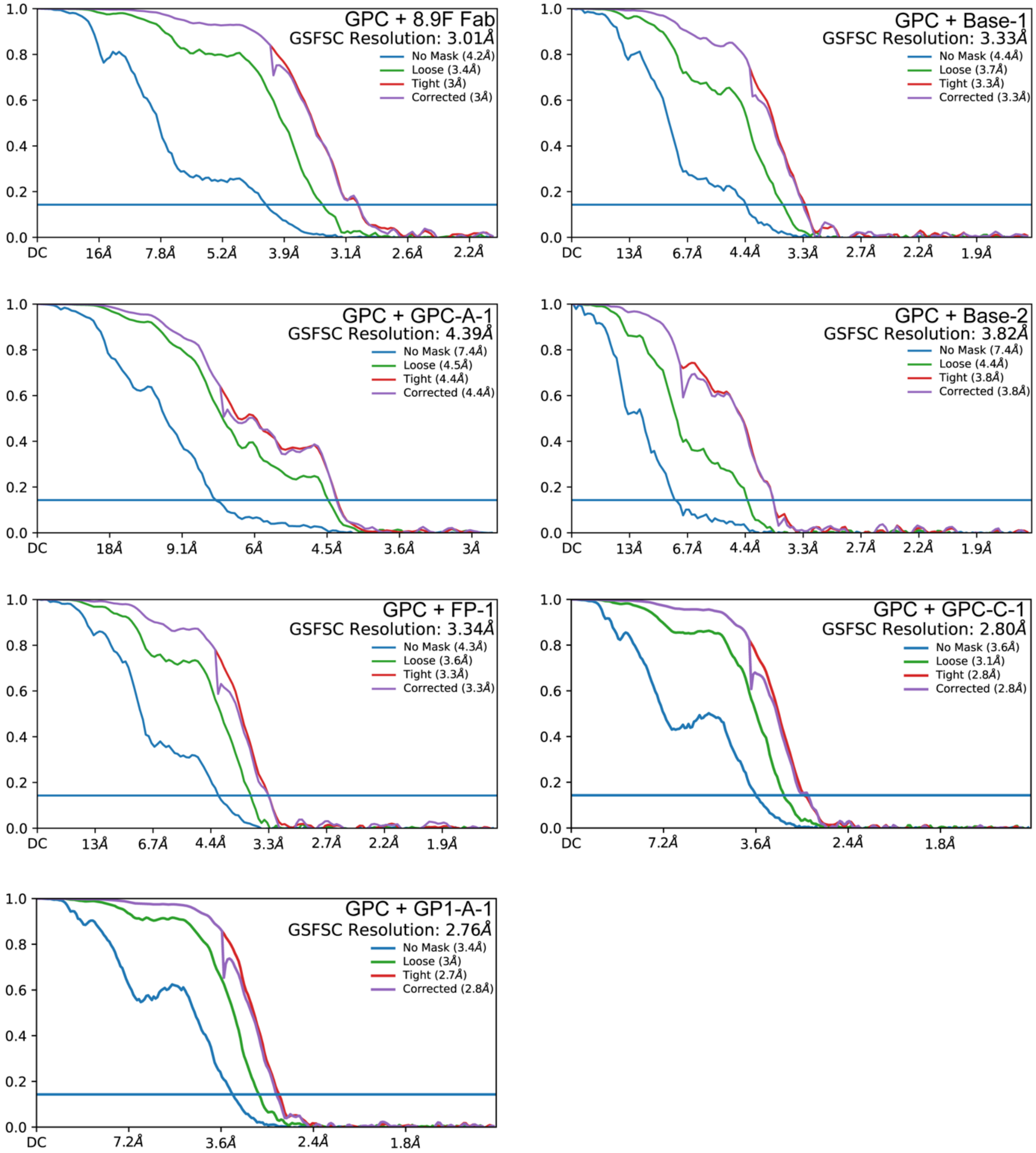


**Fig. S4: FSC plots for cryo-EM maps, related to figures 3, 4, and 6.** Reported resolutions coincide with an FSC cutoff of 0.143. Plots generated using cryoSPARC 3.2.[2]

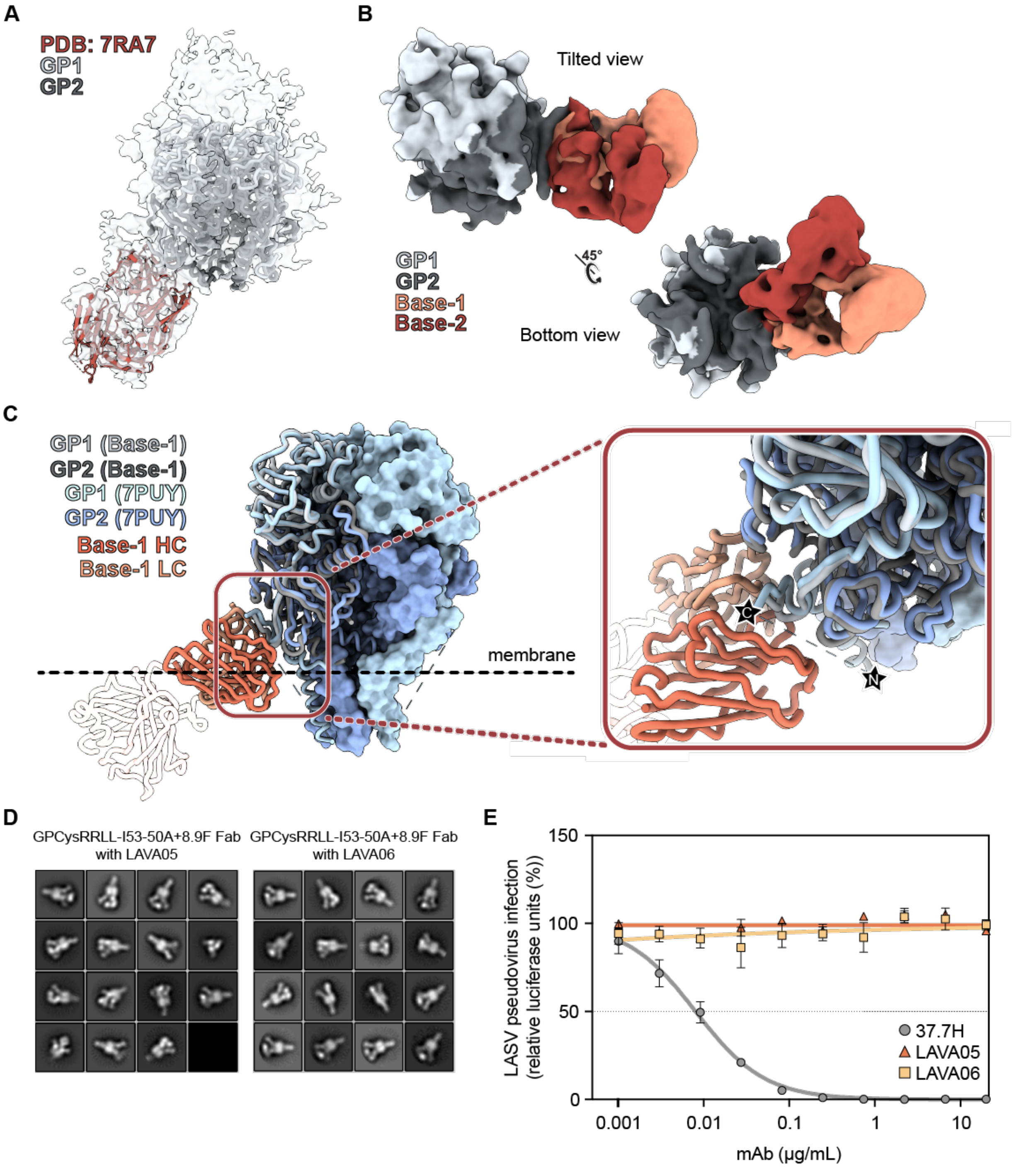


**Fig. S5. Structural differences between Base-1 and Base-2 responses, their inability to bind membrane-embedded GPC and neutralization of LAVA05 and LAVA06, related to figure 3.** **(A)** Cryo-EM map of Base-2 with a rabbit Fab (PDB: 7RA7) docked in the pFab density. **(B)** Overlaid Gaussian-filtered maps of base antibodies demonstrating the distinct angles of approach taken by Base-1 (orange) and Base-2 (red) to engage the GPC. **(C)** Overlay of full-length GPC (PDB: 7PUY[4]) with the Base-1 model revealing accessibility issues for Base-1 binding by the membrane and/or SSP. The constant region of the Fab (transparent) was generated by docking a model of a full rabbit Fab (PDB: 7RA7) into the density and removing the variable domain. The inset shows the unmodelled part of the SSP (displayed as pseudobond), referred to by Katz et al.[4] as “hydrophobic 2” (residues 36-58), with its N terminus (right) and C terminus (left) depicted as stars.[4] It is probable that the 22 amino acid “hydrophobic 2” structure resides in this area, sterically blocking binding of base-targeting pAb responses. **(D)** 2D class averages of nsEMPEM experiments with LAVA05 and LAVA05 using GPCysRRLL-I53-50A pre-complexed with 8.9F. Classes shown are the selected classes that were selected for 3D classification, reclassified to 16 classes. **(E)** Pseudovirus neutralization of LAVA05 and LAVA06. The mAb 37.7H was used as a positive control.

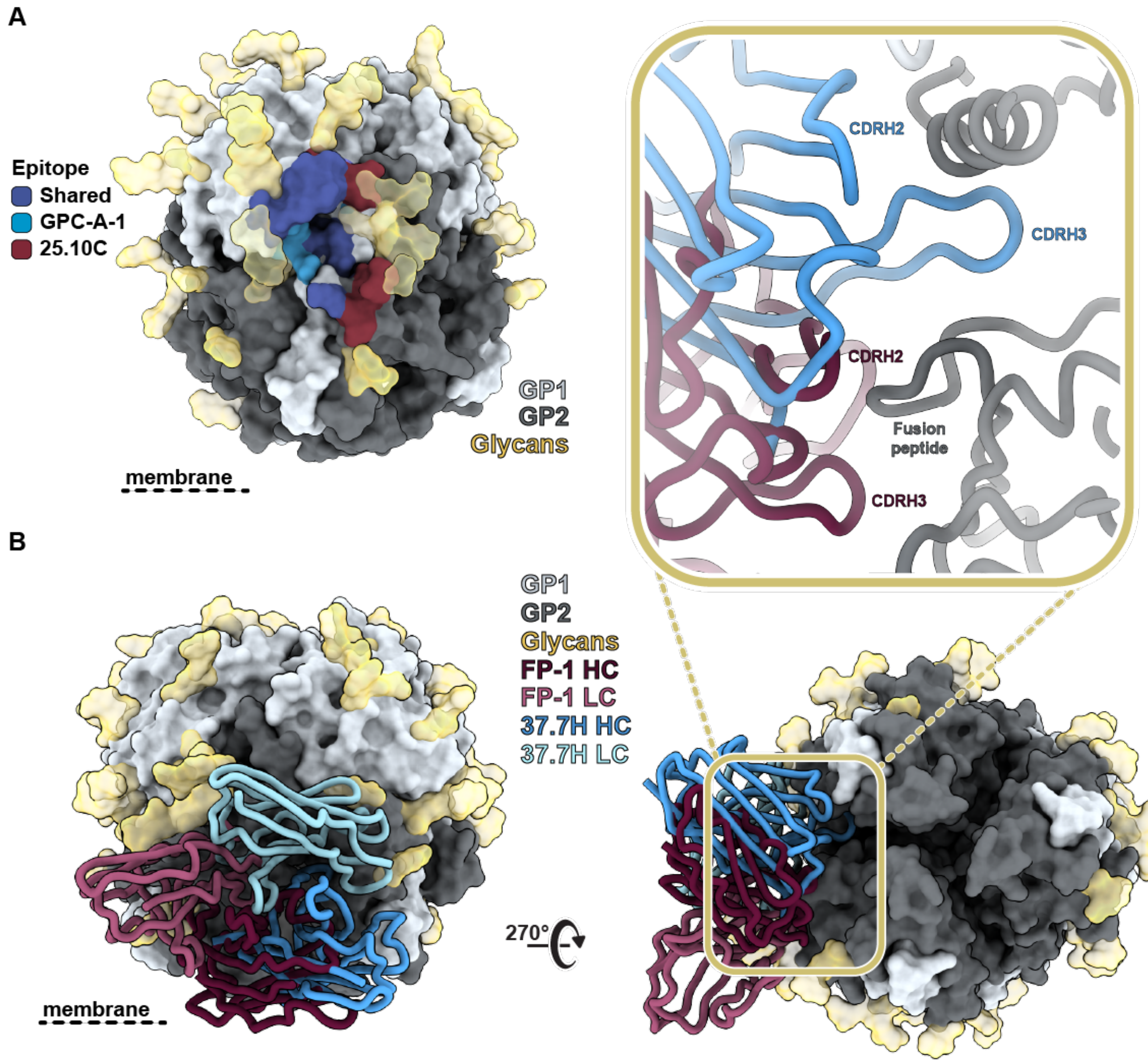


**Figure S6. Comparison of GPC-A-1 and FP-1 with previously described human NAbs, related to figure 4. (A)** Epitope footprints of GPC-A-1 and GPC-A NAb 25.10C.[5] Epitope-Analyzer[6] was used to determine (putative) contact residues using a cut-off of 4 Å between epitope and paratope for 25.10C and 6 Å for GPC-A-1 (poly-alanine model). **(B)** Comparison of GPC engagement by 37.7H and FP-1. Inset highlights the differences in fusion peptide engagement between the two antibodies.

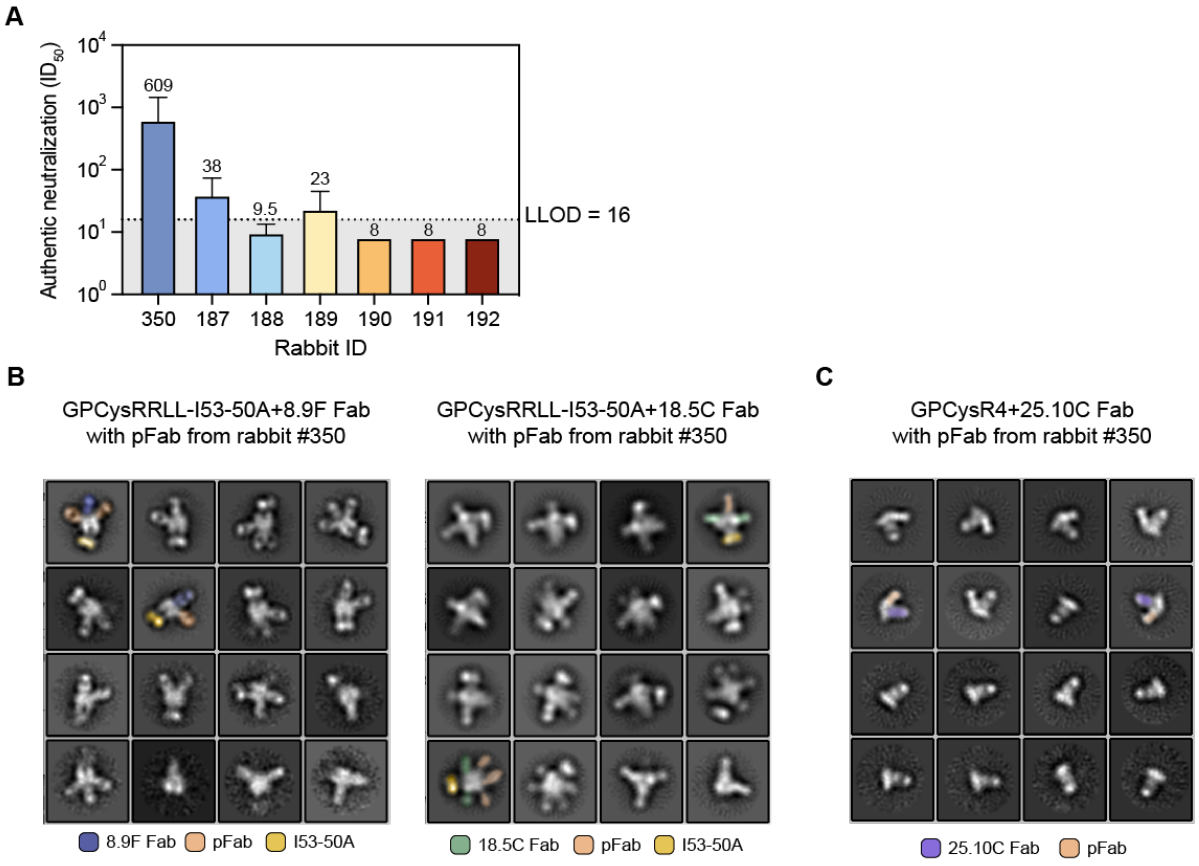


**Figure S7. Comparison of neutralization titers to authentic LASV (Josiah strain) between GPCysR4-I53-50A-vaccinated rabbits and rabbit 350 and 2D class averages of nsEMPEM experiments with pAbs from rabbit 350, related to figure 6. (A)** Autologous neutralization titers to authentic LASV from GPCysR4-I53-50A immunized rabbits (187-192) and a rabbit immunized with a GPC-derived VLP (rabbit 350). The virus neutralization titer is calculated as the geometric mean titers (GMT) of the reciprocal value of the last serum dilution at which inhibition of the cytopathic effect on infected Vero E6 cells is detectable. The initial dilution was 1:16 (LLOD) so a titer of 8 was noted when no inhibition was observed. Shown is the GMT of four technical replicates. **(B)** 2D class averages of nsEMPEM experiments with pAbs from rabbit 350 using GPCysRRLL-I53-50A pre-complexed with 8.9F (left) and 18.5C (right). Classes shown are the selected classes from two iterations of 2D classification that have been reclassified to 16 classes. Two classes are pseudo-colored according to the legend below to indicate the stabilizing Fab (8.9F Fab/18.5C Fab), pFab and I53-50A. **(C)** 2D class averages of monomeric GP nsEMPEM experiments with pAb from rabbit 350. Classes shown are the selected classes from two iterations of 2D classification that have been reclassified to 16 classes. Two classes are pseudo-colored according to the legend below to indicate the fiducial (25.10C Fab) and pFab.

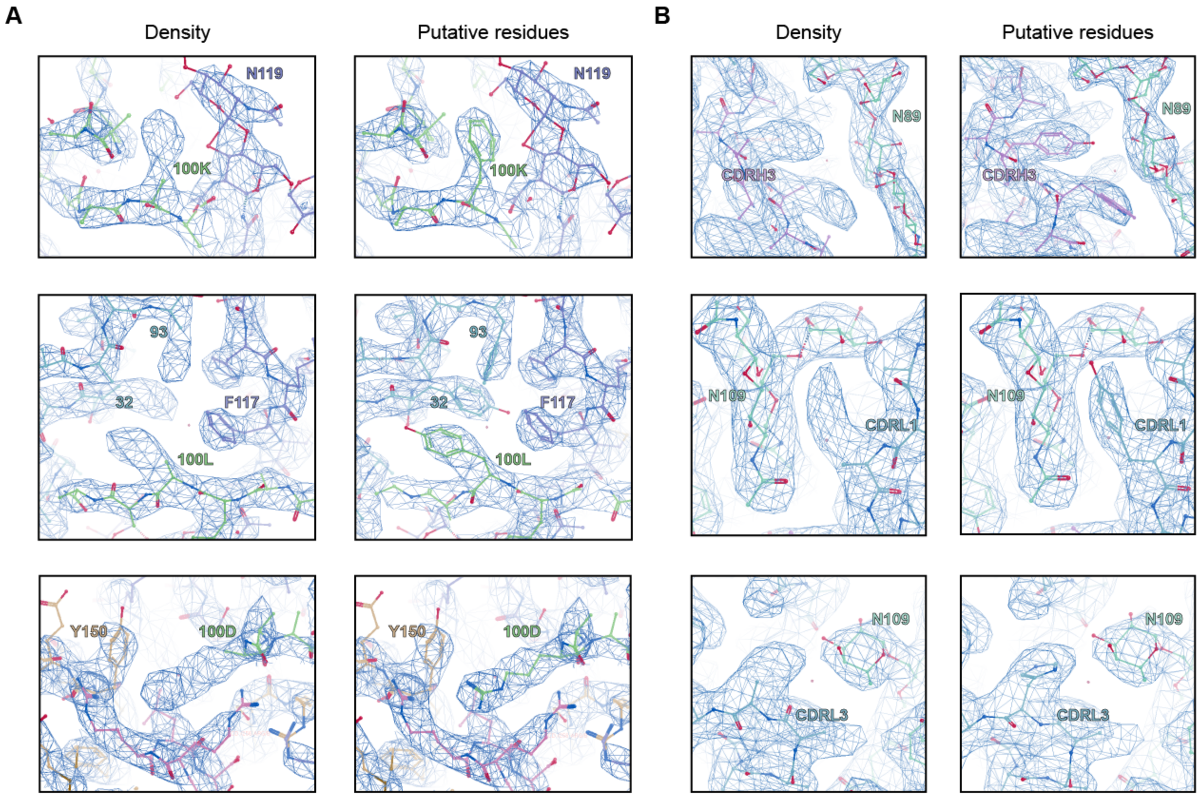


**Figure S8. Clear side-chain densities that enable speculation of GPC-pFab interactions for GPC-C-1 (A) and GP1-A-1 (B), related to figure 6. (A)** Aromatic density at position 100K in the CDRH3, making contacts with glycan N119 (top). Aromatic density at position 100L in the CDRH3, 32 in LC FR2, and and 93 in LC FR4 forming putative hydrophobic interactions with F117 in GPC (middle). Arginine density at positions 100A (not shown) and 100D in the CDRH3 presumably form cation-pi interactions with GPC's Y150 (bottom). **(B)** Aromatic densities in CDRH3 making contacts with the N89 glycan (top). Aromatic density in CDRL1 (middle) and CDRL3 (bottom) making contact with the N109 glycan.

**Table S1. 8.9F antibody interactions with GP1, related to figure 1.** Amino acid interactions at the 8.9F epitope-paratope region were determined using the online-based Epitope-Analyzer platform[6]. Glycan contacts were assessed by finding close contacts (<4 Å) of the GPC glycans with 8.9F Fab using ChimeraX[3].

| GPC Residue number | Amino acid | Chain | Atom | Fab Residue number | Amino Acid | Heavy or light chain | Distance (Å) | Interaction Type |
|---|---|---|---|---|---|---|---|---|
| 117 | PHE | A | CD1-OH | 100E | TYR | H | 4.0 | Van-der-Waals |
| 119 | ASN | A | OD1-CD1 | 100E | TYR | H | 3.7 | Van-der-Waals |
| 121 | SER | A | CB-CB | 100D | TYR | H | 3.8 | Van-der-Waals |
| 121 | SER | A | OG-N | 100D | TYR | H | 3.9 | Hydro-Bond |
| 125 | LYS | A | CE-OD2 | 100A | ASP | H | 3.9 | Van-der-Waals |
| 125 | LYS | A | NZ-OD1 | 100A | ASP | H | 3.9 | Salt-Bridge |
| 125 | LYS | A | NZ-OD2 | 100A | ASP | H | 2.8 | Salt-Bridge |
| 150 | TYR | A | CD2-O | 100H | ASP | H | 3.9 | Van-der-Waals |
| 150 | TYR | A | CE1-CG2 | 100I | VAL | H | 4.0 | Hydr-Phbc |
| 256 | ARG | A | CB-OE1 | 100L | GLN | H | 3.7 | Van-der-Waals |
| 256 | ARG | A | CZ-CE | 100G | MET | H | 3.9 | Van-der-Waals |
| 256 | ARG | A | NH2-O | 100G | MET | H | 3.2 | Hydro-Bond |
| 150 | TYR | B | OH-OD2 | 100O | ASP | H | 3.6 | Hydro-Bond |
| 256 | ARG | B | CD-CE | 100G | MET | H | 3.1 | Van-der-Waals |
| 256 | ARG | B | CD-CZ | 100C | TYR | H | 4.0 | Van-der-Waals |
| 257 | ARG | B | CB-OH | 100D | TYR | H | 3.8 | Van-der-Waals |
| 257 | ARG | B | N-OH | 100D | TYR | H | 3.7 | Hydro-Bond |
| 258 | LEU | B | N-OH | 100D | TYR | H | 3.9 | Hydro-Bond |
| 125 | LYS | C | CD-OD2 | 100O | ASP | H | 3.6 | Van-der-Waals |
| 125 | LYS | C | NZ-OD2 | 100O | ASP | H | 2.8 | Salt-Bridge |
| 150 | TYR | C | CE1-CB | 100 | ALA | H | 3.6 | Hydr-Phbc |
| 150 | TYR | C | CE2-OD1 | 100A | ASP | H | 3.4 | Van-der-Waals |
| 150 | TYR | C | CG-CE2 | 100D | TYR | H | 3.9 | Hydr-Phbc |
| 150 | TYR | C | CZ-CB | 100 | ALA | H | 3.8 | Hydr-Phbc |
| 150 | TYR | C | OH-OD1 | 100A | ASP | H | 2.6 | Hydro-Bond |
| 150 | TYR | C | OH-OD2 | 100A | ASP | H | 3.8 | Hydro-Bond |
| 256 | ARG | C | C-OD2 | 100H | ASP | H | 3.7 | Van-der-Waals |
| 256 | ARG | C | CD-CB | 100G | MET | H | 4.0 | Van-der-Waals |
| 257 | ARG | C | C-OD1 | 100H | ASP | H | 3.8 | Van-der-Waals |
| 257 | ARG | C | N-OD1 | 100H | ASP | H | 3.4 | Hydro-Bond |
| 257 | ARG | C | N-OD2 | 100H | ASP | H | 2.7 | Hydro-Bond |
| 258 | LEU | C | CA-OD1 | 100H | ASP | H | 3.8 | Van-der-Waals |
| 258 | LEU | C | N-OD1 | 100H | ASP | H | 2.9 | Hydro-Bond |
| 258 | LEU | C | N-OD2 | 100H | ASP | H | 4.0 | Hydro-Bond |
| 114 | ASN | C | C-CA | 30 | GLY | L | 3.7 | Van-der-Waals |
| 114 | ASN | C | ND2-CB | 30 | ILE | L | 3.9 | Van-der-Waals |
| 114 | ASN | C | O-N | 30 | GLY | L | 3.8 | Hydro-Bond |
| 115 | HIS | C | CB-OD1 | 31 | ASN | L | 3.9 | Van-der-Waals |
| 116 | LYS | C | N-CB | 93 | SER | L | 4.0 | Van-der-Waals |
| 117 | PHE | C | CE2-CG | 31 | ASN | L | 3.8 | Van-der-Waals |
| 117 | PHE | C | CZ-CA | 93 | SER | L | 3.9 | Van-der-Waals |
| 117 | PHE | C | CZ-CD2 | 91 | TYR | L | 3.9 | Hydr-Phbc |
| 117 | PHE | C | CZ-CE2 | 32 | PHE | L | 3.9 | Hydr-Phbc |
| 117 | PHE | C | CZ-CE2 | 91 | TYR | L | 3.9 | Hydr-Phbc |
| N119 glycan, O4-C1 GlcNAc | | A | N2-O | 54 | SER | H | 3.0 | - |
| N119 glycan, O4-C1 GlcNAc | | A | C8-O | 54 | SER | H | 3.7 | - |
| N119 glycan, O6-C1 Fuc | | A | O5-OG | 54 | SER | H | 3.7 | - |
| N119 glycan, O6-C1 Fuc | | A | C1-O | 54 | SER | H | 3.5 | - |
| N119 glycan, O4-C1 GlcNAc | | A | C7-O | 54 | SER | H | 3.8 | - |
| N119 glycan, O4-C1 GlcNAc | | A | N2-C | 54 | SER | H | 3.9 | - |
| N119 glycan, O4-C1 GlcNAc | | A | C2-O | 54 | SER | H | 4.0 | - |
| N119 glycan, O4-C1 GlcNAc | | A | C8-CA | 54 | SER | H | 4.0 | - |
| N119 glycan, O6-C1 Fuc | | A | O5-CB | 54 | SER | H | 4.0 | - |

| | | | | | | | |
|---|---|---|---|---|---|---|---|
| N119 glycan, O6-C1 Fuc | A | O4-CB | 56 | SER | H | 3.2 | - |
| N119 glycan, O6-C1 Fuc | A | O5-CB | 56 | SER | H | 3.9 | - |
| N119 glycan, O6-C1 Fuc | A | O4-CA | 56 | SER | H | 4.0 | - |
| N119 glycan, O6-C1 Man | A | O6-OG1 | 73 | THR | H | 2.9 | - |
| N119 glycan, O6-C1 Man | A | O6-CB | 73 | THR | H | 3.9 | - |
| N119 glycan, O6-C1 Man | A | O6-N | 73 | THR | H | 3.4 | - |
| N119 glycan, O6-C1 Man | A | C6-OG1 | 73 | THR | H | 3.6 | - |
| N119 glycan, O6-C1 Fuc | A | C6-O | 100A | ASP | H | 3.6 | - |
| N119 glycan, O6-C1 Fuc | A | C6-OD1 | 100B | ASP | H | 3.7 | - |
| N119 glycan, ND1-C1 GlcNAc | A | C6-CZ | 100E | TYR | H | 3.8 | - |
| N119 glycan, O4-C1 GlcNAc | A | C8-OH | 100E | TYR | H | 3.5 | - |
| N119 glycan, ND1-C1 GlcNAc | A | C5-CE1 | 100E | TYR | H | 3.7 | - |
| N119 glycan, ND1-C1 GlcNAc | A | C6-CE1 | 100E | TYR | H | 3.8 | - |
| N119 glycan, ND1-C1 GlcNAc | A | O5-CE1 | 100E | TYR | H | 3.8 | - |
| N119 glycan, ND1-C1 GlcNAc | C | C8-O | 100O | ASP | H | 3.9 | - |
| N119 glycan, ND1-C1 GlcNAc | C | C8-CD2 | 100Q | TYR | H | 3.8 | - |
| N119 glycan, ND1-C1 GlcNAc | C | O7-OH | 100S | TYR | H | 3.2 | - |
| N119 glycan, ND1-C1 GlcNAc | C | O7-CZ | 100S | TYR | H | 3.8 | - |
| N119 glycan, ND1-C1 GlcNAc | C | O7-CE2 | 100S | TYR | H | 3.5 | - |
| N119 glycan, O6-C1 Fuc | C | O4-OD1 | 31 | ASN | L | 2.8 | - |
| N119 glycan, O6-C1 Fuc | C | C4-OD1 | 31 | ASN | L | 3.7 | - |
| N119 glycan, O6-C1 Fuc | C | O4-CG | 31 | ASN | L | 3.7 | - |
| N119 glycan, O6-C1 Fuc | C | O3-C | 31 | ASN | L | 3.7 | - |
| N119 glycan, O6-C1 Fuc | C | O3-CA | 31 | ASN | L | 3.7 | - |
| N119 glycan, O6-C1 Fuc | C | O4-CB | 31 | ASN | L | 3.8 | - |
| N119 glycan, O6-C1 Fuc | C | C4-CB | 31 | ASN | L | 3.9 | - |
| N119 glycan, O6-C1 Fuc | C | O3-N | 32 | PHE | L | 2.8 | - |
| N119 glycan, O6-C1 Fuc | C | O3-CA | 32 | PHE | L | 3.7 | - |
| N119 glycan, O6-C1 Fuc | C | C5-CZ | 32 | PHE | L | 3.8 | - |
| N119 glycan, O6-C1 Fuc | C | C4-CE1 | 32 | PHE | L | 3.8 | - |
| N119 glycan, O6-C1 Fuc | C | C4-CZ | 32 | PHE | L | 3.9 | - |
| N119 glycan, O6-C1 Fuc | C | C3-CD1 | 32 | PHE | L | 3.9 | - |
| N119 glycan, O6-C1 Fuc | C | C3-CE1 | 32 | PHE | L | 3.9 | - |
| N119 glycan, O6-C1 Fuc | C | C3-N | 32 | PHE | L | 3.8 | - |
| N119 glycan, O6-C1 Fuc | C | C5-CE1 | 32 | PHE | L | 3.9 | - |
| N119 glycan, O6-C1 Fuc | C | C6-CZ | 32 | PHE | L | 4.0 | - |
| N119 glycan, ND1-C1 GlcNAc | C | O3-OE2 | 50 | GLU | L | 3.5 | - |
| N119 glycan, ND1-C1 GlcNAc | C | O3-OE1 | 50 | GLU | L | 4.0 | - |
| N119, O4-C1 GlcNAc | C | C1-OE1 | 50 | GLU | L | 3.7 | - |
| N119 glycan, ND1-C1 GlcNAc | C | O3-CD | 50 | GLU | L | 3.8 | - |
| N119, O6-C1 Man | C | O6-CG1 | 51 | VAL | L | 3.5 | - |
| N119, O6-C1 Man | C | O6-CB | 51 | VAL | L | 3.9 | - |
| N119, O4-C1 Man | C | O5-O | 51 | VAL | L | 4.0 | - |
| N119, O4-C1 Man | C | O2-CB | 53 | SER | L | 3.4 | - |
| N119, O4-C1 Man | C | O2-OG | 53 | SER | L | 3.8 | - |
| N119, O3-C1 Man | C | O3-OG | 53 | SER | L | 3.2 | - |
| N119, O4-C1 Man | C | O2-CA | 53 | SER | L | 3.8 | - |
| N119 glycan, O6-C1 Fuc | C | O3-CE | 66 | LYS | L | 3.6 | - |
| N119 glycan, O6-C1 Fuc | C | O3-CD | 66 | LYS | L | 3.8 | - |
| N119 glycan, O6-C1 Fuc | C | O3-NZ | 66 | LYS | L | 3.9 | - |

**Table S2: Electron Microscopy Data Bank deposition information, related to figures 2 and 3.** Maps are accessible at emdataresource.org using the listed codes. Additional maps can be found on the “Download” tab of each entry.

| | EMDB ID | Time point | Specificity | # particles (composite) | Map | Microscope | Pixel size |
|---|---|---|---|---|---|---|---|
| Rabbit 187 | EMD-43174 | 30 | Base | 7,212 | additional map | Tecnai Spirit | 2.06 |
| | EMD-43174 | 30 | FP | 10,038 | additional map | Tecnai Spirit | 2.06 |
| | EMD-43174 | 30 | GPC-A | 16,689 | main map | Tecnai Spirit | 2.06 |
| | EMD-43175 | 18 | Base | 8,882 | additional map | Tecnai Spirit | 2.06 |
| | EMD-43175 | 18 | FP | 8,905 | additional map | Tecnai Spirit | 2.06 |
| | EMD-43175 | 18 | GPC-A | 12,945 | main map | Tecnai Spirit | 2.06 |
| Rabbit 188 | EMD-43176 | 30 | Base | 9,484 | main map | Tecnai Spirit | 2.06 |
| Rabbit 189 | EMD-43177 | 30 | Base | 12,037 | additional map | Tecnai Spirit | 2.06 |
| | EMD-43177 | 30 | GPC-A | 13,250 | main map | Tecnai Spirit | 2.06 |
| | EMD-43178 | 18 | Base | 5,061 | additional map | Tecnai Spirit | 2.06 |
| | EMD-43178 | 18 | GPC-A | 2,937 | main map | Tecnai Spirit | 2.06 |
| Rabbit 190 | EMD-43179 | 30 | Base | 6,805 | additional map | Tecnai Spirit | 2.06 |
| | EMD-43179 | 30 | FP | 11,371 | main map | Tecnai Spirit | 2.06 |
| Rabbit 191 | EMD-43180 | 30 | Base | 11,393 | main map | Tecnai Spirit | 2.06 |
| Rabbit 192 | EMD-43181 | 30 | Base | 15,413 | additional map | Tecnai Spirit | 2.06 |
| | EMD-43181 | 30 | FP | 15,136 | main map | Tecnai Spirit | 2.06 |
| LAVA05 | EMD-43182 | 30 | Base | 17,515 | main map | Tecnai Spirit | 2.06 |
| LAVA06 | EMD-43183 | 30 | Base | 19,440 | main map | Tecnai Spirit | 2.06 |

**Table S3: Cryo-EM data collection and atomic model refinement, related to figures 3, 4, 5, and 6.**

| | GPC bound to mAb 8.9F | GPC bound to pAb GPC-A-1 | GPC bound to pAb Base-1 | GPC bound to pAb Base-2 | GPC bound to pAb FP-1 | GP monomer bound to INT-1 and INT-2 | GPC bound to pAb GPC-C-1 | GPC bound to pAb GP1-A-1 |
|---|---|---|---|---|---|---|---|---|
| **Access codes** | | | | | | | | |
| PDB | 8ER3 | 8TYB | 8TYC | NA | 8TYE | N/A | 8VCV | 8VE8 |
| EMDB | EMD-28549 | EMD-41712 | EMD-41713 | EMD-41714 | EMD-41715 | EMD-41716 | EMD-43141 | EMD-43168 |
| Genbank | NP_694870.1 | NP_694870.1 | NP_694870.1 | NP_694870.1 | NP_694870.1 | NP_694870.1 | NP_694870.1 | NP_694870.1 |
| **Data collection and processing** | | | | | | | | |
| Microscope | Titan Krios | Titan Krios | Titan Krios | Titan Krios | Titan Krios | TFS Glacios | TFS Glacios II | TFS Glacios II |
| Magnification | 130,000 | 130,000 | 130,000 | 130,000 | 130,000 | 190000 | 190000 | 190000 |
| Voltage (kV) | 300 | 300 | 300 | 300 | 300 | 200 | 200 | 200 |
| Electron exposure ($e^-/Å^2$) | 50.0 | 49.1 | 50.35 | 50.35 | 50.35 | 49.5 | 40.7 | 40.7 |
| Defocus range (μm) | -0.7 to -2.0 | -0.7 to -2.0 | -0.7 to -2.0 | -0.7 to -2.0 | -0.7 to -2.0 | -0.8 to -1.4 | -0.8 to -2.0 | -0.8 to -2.0 |
| Pixel size (Å) | 1.045 | 0.725 | 0.833 | 0.833 | 0.833 | 0.725 | 0.718 | 0.718 |
| Imposed Symmetry | C1 | C1 | C1 | C1 | C1 | C1 | C1 | C1 |
| Final particle number | 73,222 | 30,182 (symmetry expanded) | 61,338 (symmetry expanded) | 14,690 (symmetry expanded) | 77,295 (symmetry expanded) | 20,839 | 89,195 | 146,722 |
| Map resolution (Å) | 3.0 | 4.4 | 3.3 | 3.8 | 3.3 | 8.1 | 2.8 | 2.8 |
| FSC Threshold | 0.143 | 0.143 | 0.143 | 0.143 | 0.143 | 0.143 | 0.143 | 0.143 |
| Map sharpening B-factor ($Å^2$) | -35 | -96 | -60 | -80 | -90 | -896 | -58 | -64 |
| **Model refinement and validation** | | | | | | | | |
| Total Residues | 1389 | 1369 | 1353 | N/A | 1351 | N/A | 1343 | 1329 |
| Amino-acids | 1300 | 1281 | 1273 | | 1266 | | 1267 | 1260 |
| Carbohydrates | 89 | 88 | 80 | | 85 | | 76 | 69 |
| RMSD Bonds | 0.02 | 0.02 | 0.02 | | 0.02 | | 0.02 | 0.02 |
| RMSD Angles | 1.7 | 1.8 | 1.7 | | 1.8 | | 1.7 | 1.8 |
| **Ramachandran** | | | | | | | | |
| Outliers (%) | 0 | 0 | 0 | | 0 | | 0 | 0 |
| Allowed (%) | 2.12 | 4.38 | 4.48 | | 3.79 | | 2.89 | 2.75 |
| Favored (%) | 97.88 | 95.62 | 95.52 | | 96.21 | | 97.11 | 97.25 |
| Rotamer outliers (%) | 0 | 0 | 0 | | 0 | | 0 | 0 |
| Clash score | 2.1 | 3.1 | 3.1 | | 3.1 | | 3.2 | 2.7 |
| Molprobity score | 1.0 | 1.4 | 1.4 | | 1.4 | | 1.3 | 1.2 |
| FSC model (0/0.143/0.5) | 2.7/3.0/3.3 | 4.2/4.3/4.6 | 3.0/3.3/3.7 | | 3.2/3.4/3.9 | | 2.7/2.8/3.0 | 2.7/2.7/2.9 |
| EMRinger score | 4.58 | 1.42 | 2.83 | | 2.09 | | 5.5 | 5.55 |
